# The evolution of local adaptation in long-lived species

**DOI:** 10.1101/2024.07.17.603878

**Authors:** Loraine Hablützel, Charles Mullon, Max Schmid

## Abstract

Many species experience heterogeneous environments and adapt genetically to local conditions. The extent of such local adaptation has been shown to depend on a balance between divergent selection and gene flow, but also on other factors such as phenotypic plasticity or the genetic architecture of traits. Here, we explore the role of life history in this process. We develop a quantitative genetics model and run individual-based simulations to contrast the evolution of local adaptation between short- and long-lived species. We show that life history does not have a unidirectional effect on local adaptation. Instead, local adaptation varies with a species’ life cycle and how this cycle modulates the scheduling of selection and dispersal among stage classes. When a longer generation time is associated with more frequent events of selection than dispersal, local adaptation is more pronounced in long-lived than in short-lived species. If otherwise dispersal occurs more frequently than selection, long-lived species evolve weaker local adaptation. Our simulations confirm these findings and further highlight how the scheduling of selection and dispersal shape additive genetic variance, effective dispersal between patches, and the genetic response at quantitative trait loci. Taken together, our results suggest that the effect of longevity on local adaptation depends on the specifics of a species’ life cycle, potentially explaining why current meta-analyses have not consistently detected this effect.

## 1 Introduction

Most species experience a heterogeneous environment, seeing variation in biotic and/or abiotic conditions across their range. Such spatial heterogeneity can lead species to adapt genetically to local conditions such that local genotypes achieve higher fitness than genotypes from other habitats (Kawecki, 2008; Blanquart et al., 2013). This fitness pattern, which is referred to as local adaptation, has been described in a wide range of taxa and spatial scales (Leimu and Fischer, 2008; Hereford, 2009; Halbritter et al., 2018). In Atlantic salmon (*Salmo salar*), for instance, local female fish show a nine-fold greater reproductive success compared to females from other sites of the same river system (Mobley et al., 2019). Genotypes of Thale cress (*Arabidopsis thaliana*) at the southern range edge in Italy have fitness that is 14-fold greater than genotypes originating from the northern range edge in Sweden (Postma and Ågren, 2016). Accordingly, understanding the evolution of local adaptation through genetic divergence between habitats has been an important goal within evolutionary biology (Rellstab et al., 2015; Lind et al., 2018), as well as for more applied sciences such forestry, fishery, or nature conservation (Aitken et al., 2008; Fraser et al., 2011; Kuparinen et al., 2010).

Theoretical models have proven useful to understand evolution in a spatial context and therefore local adaptation. One important and well-established notion is the limiting effect of gene flow on local adaptation, as dispersal hinders genetic differentiation between localities (Haldane, 1948; Slatkin, 1973; Felsenstein, 1976; Lenormand, 2002). Because gene flow is ubiquitous, genetic divergence is most often thought to be incomplete, reflecting a balance between migration and selection that leads to less than optimal divergence (Hendry et al., 2001). The equilibrium level of genetic divergence may also depend on many other factors, such as asymmetry in gene flow (Kawecki, 2008; Sexton et al., 2009; Polechová and Barton, 2015), phenotypic plasticity (Crispo, 2008; Thibert-Plante and Hendry, 2011; Schmid and Guillaume, 2017; Soularue et al., 2023), ecological interactions within and between species (Connallon, 2015; Bolnick and Stutz, 2017; Schmid et al., 2024), or the genetic architecture of traits (Bulmer, 1972; Guillaume, 2011; Yeaman, 2015; Pontz and Bürger, 2022). These various factors influence local adaptation through multiple pathways, e.g. by modulating the strength of divergent selection acting on the focal trait, modifying the extent of gene flow, or affecting the evolutionary response to selection.

Most of this theory on the evolution of local adaptation is built for annual semelparous species with simple life histories (most often modelled as a Wright-Fisher process, e.g. Hendry et al., 2001; Thibert-Plante and Hendry, 2011; Guillaume, 2011; Schmid and Guillaume, 2017; Pontz and Bürger, 2022; Soularue et al., 2023). But many organisms are long-lived and follow more complicated life cycles, transitioning through various stages, e.g. from resting to dispersal stages or from pre-mature to reproductive stages (Salguero-Gómez et al., 2017; Healy et al., 2019). The implications of longevity and stage structure for the evolution of local adaptation are not straightforward. Stage classes could differ from each other in their dispersal capability and in their exposure to selection, such that local adaptation might evolve differently in each stage. In addition, individuals might stay in a stage for several years or transit to other classes, thus modulating genetic variance in each stage and therefore the response to selection (e.g., Lande, 1982; Barfield et al., 2011). While these observations point to a potentially important role for longevity and stage structure, meta-analyses have so far not found any systematic effect for these in shaping local adaptation (Leimu and Fischer, 2008; Halbritter et al., 2018; Runquist et al., 2020). Specifically, current evidence suggests that local adaptation does not vary significantly between life-history types (e.g., annual vs perennial plants) across systems.

Here, we investigate the evolution of local adaptation in long-lived species using a quantitative genetics model and offer potential resolution to these inconsistencies. We model the evolution of a single polygenic trait for a species with stage-structured life history (as in Barfield et al., 2011), and incorporate stage-specific dispersal and viability selection. We find that the evolution of local adaptation varies with a species’ life history, but not in a systematic manner. Rather, our results reveal that greater generation time and iteroparity can either increase or decrease genetic divergence between patches, and that this depends on the scheduling of dispersal and selection throughout life.

## 2 Model

We consider a monoecious diploid species that is divided between two large patches connected by dispersal (Fig. 1a). These patches show different environments such that different values of a single evolving trait (e.g. root length) maximize survival in each patch (Fig. 1b). Individuals follow a stage-structured life history, with a total of *k* stage classes. Each stage *s* ϵ {1,…, *k*} might represent an age, size, developmental, or social class (Fig. 1c). Individuals belonging to different stage classes may differ in their dispersal proclivity, in their exposure to selection, and in their demographic contribution to other classes, e.g. through maturation, stage progression, or reproduction.

**Figure 1:**
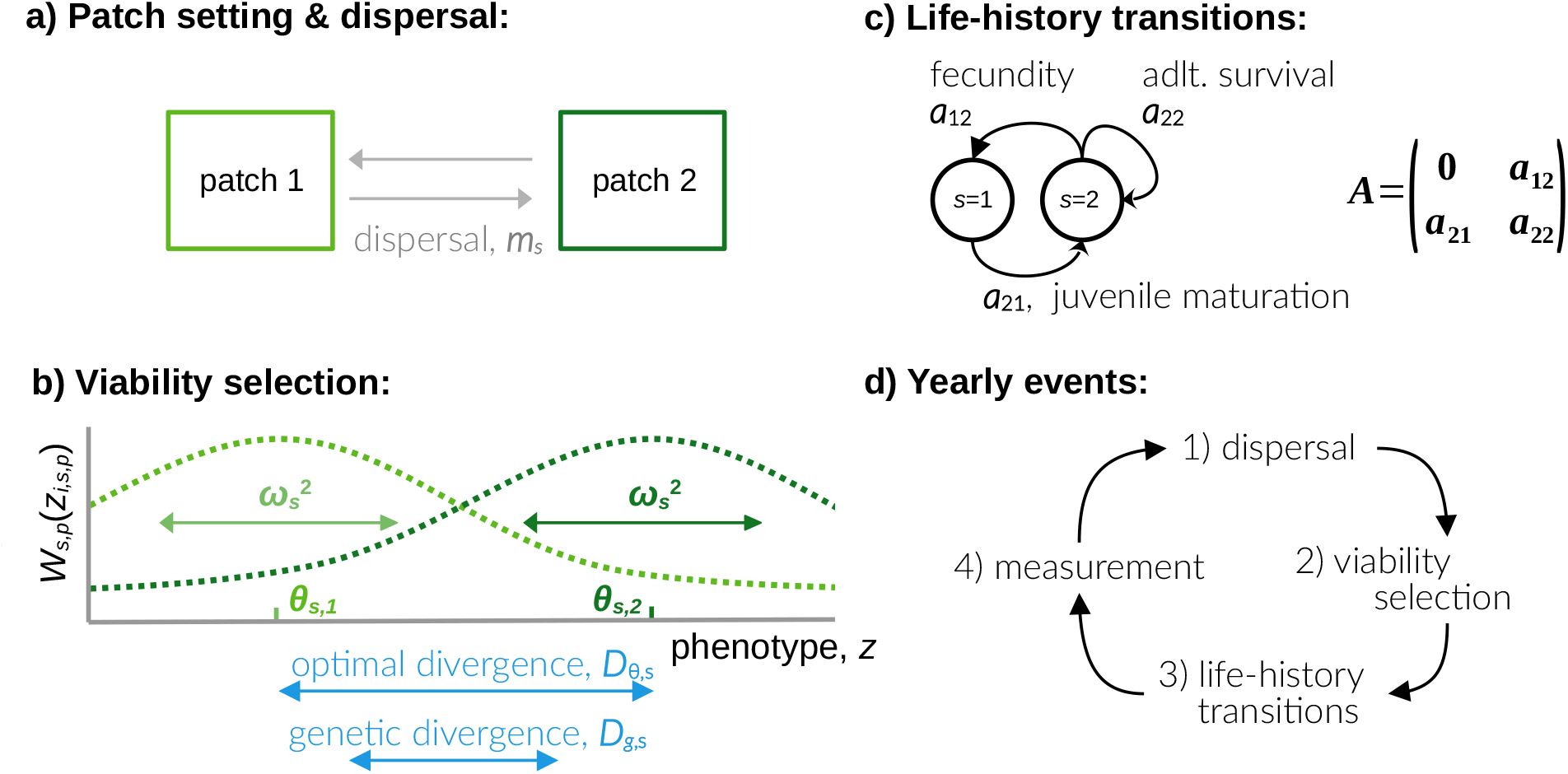
We model a single species in a two-patch setting (a). Individuals disperse with a stage-specific dispersal probability *m*_*s*_ and survive viability selection with probability *W*_*s,p*_ (*z*_*i,s,p*_) (b). Survival follows a Gaussian function and varies with individual trait value *z*, stage- and patch-specific phenotypic optima *θ*_*s,p*_, and the width of the survival function 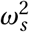.The focal species has a two-stage life history with short-lived juveniles and long-lived adults (c). Note, however, that our mathematical analysis can also be applied to other life-history strategies. Within each year, individuals first disperse, then experience selection and commit stage transitions, before we quantify local adaptation (d).

### 2.1 Life-cycle events

We assume that within each year, the following events happen in sequence: 1) dispersal; 2) viability selection; 3) life-history transitions; 4) census (Fig. 1d; we consider alternative sequences in Appendix A).

#### Dispersal

In each patch, individuals belonging to class *s* disperse to the other patch with probability *m*_*s*_ and stay put with probability 1 − *m*_*s*_. The per-year dispersal probability is thus the same in both patches but can differ among stage classes. To simplify notation, we denote the annual dispersal propensity as 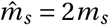. According to this rescaling, random dispersal in stage *s* occurs when 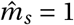 (i.e. *m*_*s*_ *=* 0.5), such that after this life-cycle event a randomly sampled individual in stage *s* is equally likely to be in either patch.

#### Selection

Each individual *i* in class *s* in patch *p* survives with a probability *W*_*s,p*_ (*z*_*i,s,p*_) that depends on its trait value *z*_*i,s,p*_ such that

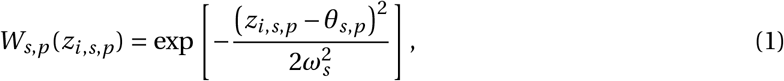

where *θ*_*s,p*_ is the trait value that maximizes survival in stage *s* in patch *p*, and 1/*ω*^2^ is the strength of selection in class *s* (i.e. it controls the rate at which survival declines with the distance between *z*_*i,s,p*_ and *θ*_*s,p*_; Fig. 1b, green dotted lines). Selection thus favors different trait values in each patch when *θ*_*s*,1_ ≠ *θ*_*s*,2_. For instance, plants might need a greater frost tolerance in one patch compared to the other to maximize survival. We denote the divergence in phenotypic optima between patches in class *s* by 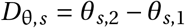 (Fig. 1b). Such divergence gives rise to divergent selection, favoring the evolution of phenotypic differentiation among patches, and thus local adaptation.

#### Life-history transitions

From conception to death, individuals pass through a series of stage classes, with at most one transition occurring per individual per year. Such life-history transition can involve various demographic processes, such as sexual reproduction, stage progression or regression. These processes are modelled as a matrix population model, captured by a *k×k* matrix **A** whose entry *a*_*sr*_ gives the expected contribution to class *s* that an individual in stage *r* makes (e.g. Fig. 1c). That is, if we denote by *n*_*s,p*_ the density of individuals in class *s* in patch *p* in a given year, and collect these densities in the column vector **n**_*p*_ *=* (*n*_1,*p*_, …, *n*_*s,p*_)^T^ (where uppercase T means transpose), densities after life-history transitions are given by 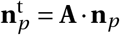 (where t denotes variables after life-history transitions such that 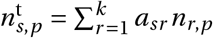. We assume that life-history transitions occur in the same way in each patch (i.e. transition rates *a*_*sr*_ are independent of *p*) and that they are constant over time (i.e. *a*_*sr*_ are parameters in our model).

### 2.2 Analyses

We investigate the evolution of a single quantitative genetic trait *z* ∈ **ℝ** in this species. The trait value *z*_*i,s,p*_ of individual *i* in stage *s* and patch *p* is given by *z*_*i,s,p*_ *= g*_*i,s,p*_ *+ e*_*i,s,p*_, where *g*_*i,s,p*_ is an additive genetic effect (i.e. the breeding value), and *e*_*i,s,p*_ is a random environmental effect which is assumed to be determined at birth (normally distributed with mean 0 and variance *σ*^2^). We quantify local adaptation as the average genetic divergence between patches for each stage *s*: 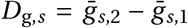, where 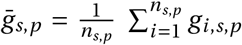 is the average additive genetic value in patch *p* among individuals in stage *s*. Let us denote genetic divergence at evolutionary equilibrium as 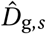,where the effects of dispersal, selection, and life-history transition balance. We use two approaches to investigate 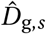.

First, we derive an evolutionary quantitative genetics model; we apply the general approach of Lande (1982) and Barfield et al. (2011), who consider evolution under age- and stage-structure respectively, to investigate the evolution of local adaptation as in Hendry et al. (2001), who tackles local adaption for a semelparous species (Appendix A for details). This approach assumes that the population size is large and that (infinitely) many unlinked autosomal loci with small additive effects contribute to the trait. Genetic and phenotypic values (i.e. *g*_*i,s,p*_ and *z*_*i,s,p*_) are normally distributed within each patch and stage and the genetic variance, 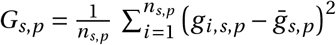, and phenotypic variance, 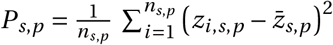 in all stages *s* are constant over time and equal among patches (i.e. *G*_*s,p*_ = *G*_*s*_ and *P*_*s,p*_ *= P*_*s*_ for all *p*).

Second, to relax the assumptions of this quantitative genetics model, we run stochastic individual-based simulations using Nemo-Age (Cotto et al., 2020, Appendix B). These simulations allow us to model a finite population size and having a finite number of quantitative loci contributing to the trait. As a baseline, we assume that each patch carries a maximum of 10,000 individuals, and that the trait is underlain by 50 unlinked loci with additive effects that evolve according to a continuum-of-alleles model (see Appendix B for more details). We use these simulations to track the evolution of genetic divergence, as well as investigate the life-cycle effects on additive genetic variance *G*_*s,p*_ and effective dispersal 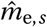 (i.e. the backward migration probability, which is the probability that a randomly sampled individual in stage *s* during census is not in its natal patch).

For ease of presentation, we focus in the main text on a species with two stage classes (*k =* 2) that is at demographic equilibrium (i.e. when the population growth rate is *λ =* 1; but see Appendix A for results with arbitrary *k* and *λ*). The first stage consists of premature juveniles (*s =* 1), and the second stage consists of reproducing adults (*s =* 2). Juveniles either mature or die within a single year (with probabilities *a*_21_ and 1 − *a*_21_, respectively). In contrast, adults are potentially long-lived and survive from one year to the next with probability *a*_22_ (Fig. 1c). The generation time for this life history, that is the mean age of a parent, is

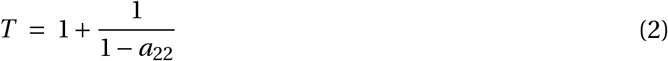

(see eqs. A49-A52 in Appendix for a derivation). As expected, greater adult survival *a*_22_ leads to a greater generation time. Note that we consider long-lived juveniles and short-lived adults, which can model plant species with a seedbank, in section A.3.4 of Appendix A.

## 3 Results

### Life-cycle events determine whether longevity is associated with local adaptation

We investigate the equilibrium genetic divergence among patches in adults 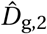 under different scenarios for the life cycle. We first consider the case where only juveniles are exposed to selection and disperse (i.e. 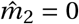 and 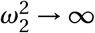). This might apply to perennial plant species with seed or pollen dispersal where local adaptation depends on a seed trait. We show in Appendix A.3 that in this case,

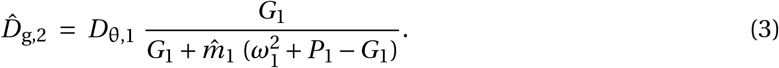

According to eq. (3), genetic divergence increases with: (i) the strength of divergent selection (i.e. with greater 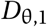 and smaller 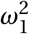); (ii) greater genetic variance *G*_1_ for the trait; and (iii) greater heritability (i.e. lower *P*_1_ −*G*_1_). Dispersal between patches 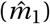 meanwhile limits local adaptation (see Fig. 2, red line). These findings reflect the common notion that local adaptation depends on the balance between divergent selection and gene flow (Lenormand, 2002). Interestingly, genetic divergence here does not change with vital rates such as adult survival (*a*_22_) or juvenile maturation (*a*_12_; see Fig. 3a, red line). In fact, eq. (3) is equal to eq. (7) in Hendry et al. (2001) (see Fig. 2, black dashed line), which considers genetic divergence in an annual (semelparous) species with dispersal followed by selection (here though with stage-specific parameters, see also our Appendix A.3.1). This indicates that a long-lived species in a scenario where only juveniles are exposed to selection and disperse faces the same balance between selection and dispersal as an annual species.

**Figure 2:**
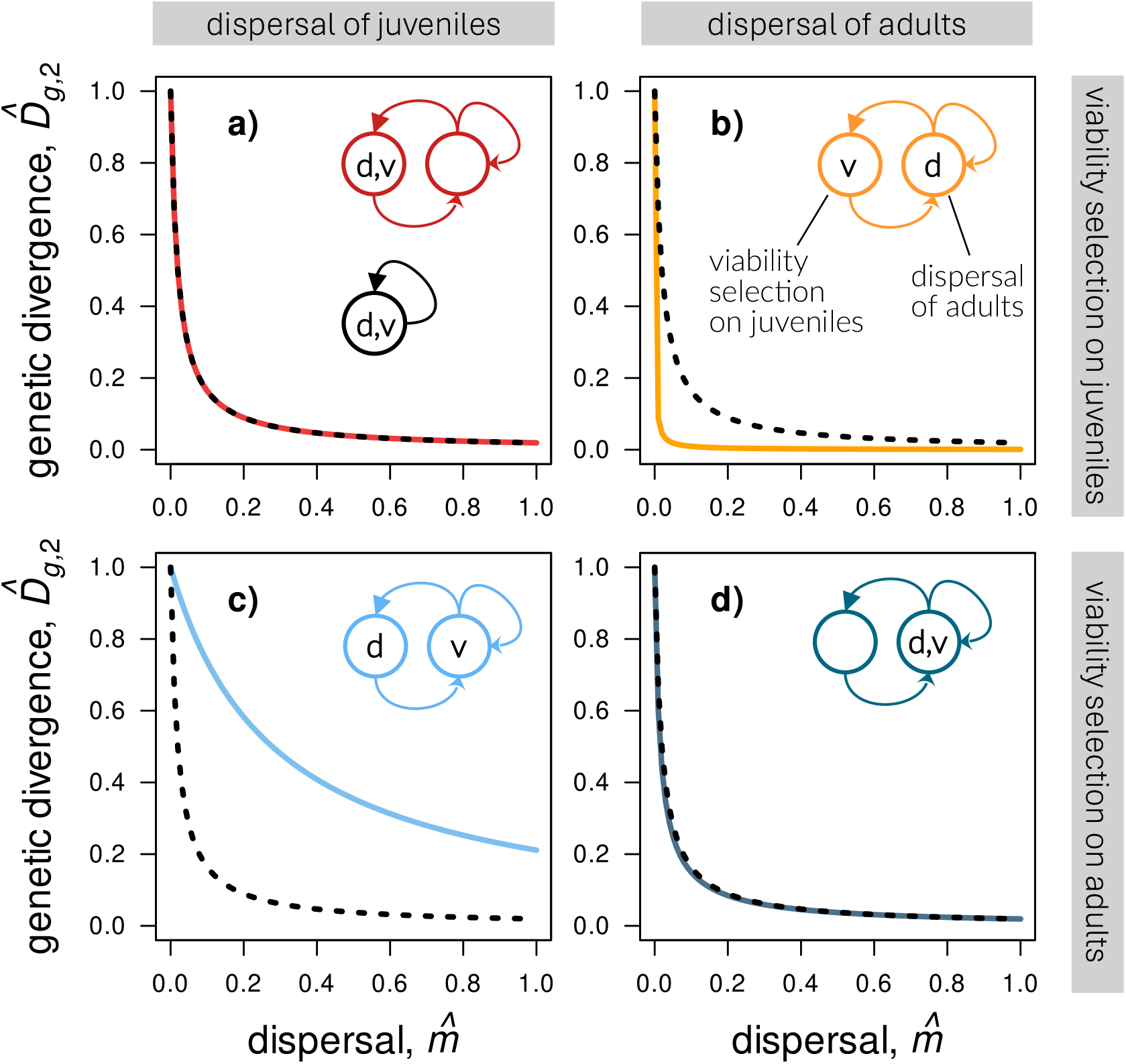
These four graphs show the extent of genetic divergence in the adult life stage for a long-lived species with four distinct timings of dispersal (as indicated by the letter *d* in the life-history diagram) and viability selection (symbolized by the letter *v*). On the left, we show species with juvenile dispersal and on the right we show species with adult dispersal. Top panels show species where viability selection acts on juveniles and bottom panels show species where viability selection acts on adults. Genetic divergence in the long-lived species is compared to local adaptation in an annual species (black dashed line) for the same ecological scenario. In graph **a)**, selection and dispersal both happen in the juvenile stage and the long-lived species evolves local adaptation like an annual species. In graph **b)**, juveniles experience viability selection and adults disperse such that the long-lived species evolves lower extents of local adaptation. In graph **c)**, juveniles disperse but adults are under selection, whereby the long-lived species evolves greater local adaptation than an annual species. In graph d), selection and dispersal act in the adult stage and the long-lived species evolves slightly lower degrees of local adaptation. Results are plotted from eqs. 3-6 with *D*_θ_ *=* 1, *ω*^2^ *=* 25, *G =* 0.5, *P =* 1, *a*_22_ *=* 0.95, *a*_11_ *=* 0, *a*_21_ *=* 0.5, and *a*_12_ *=* 0.1. As a consequence, the long-lived species has a growth rate of *λ =* 1.0 and a generation time of 21 years.

**Figure 3:**
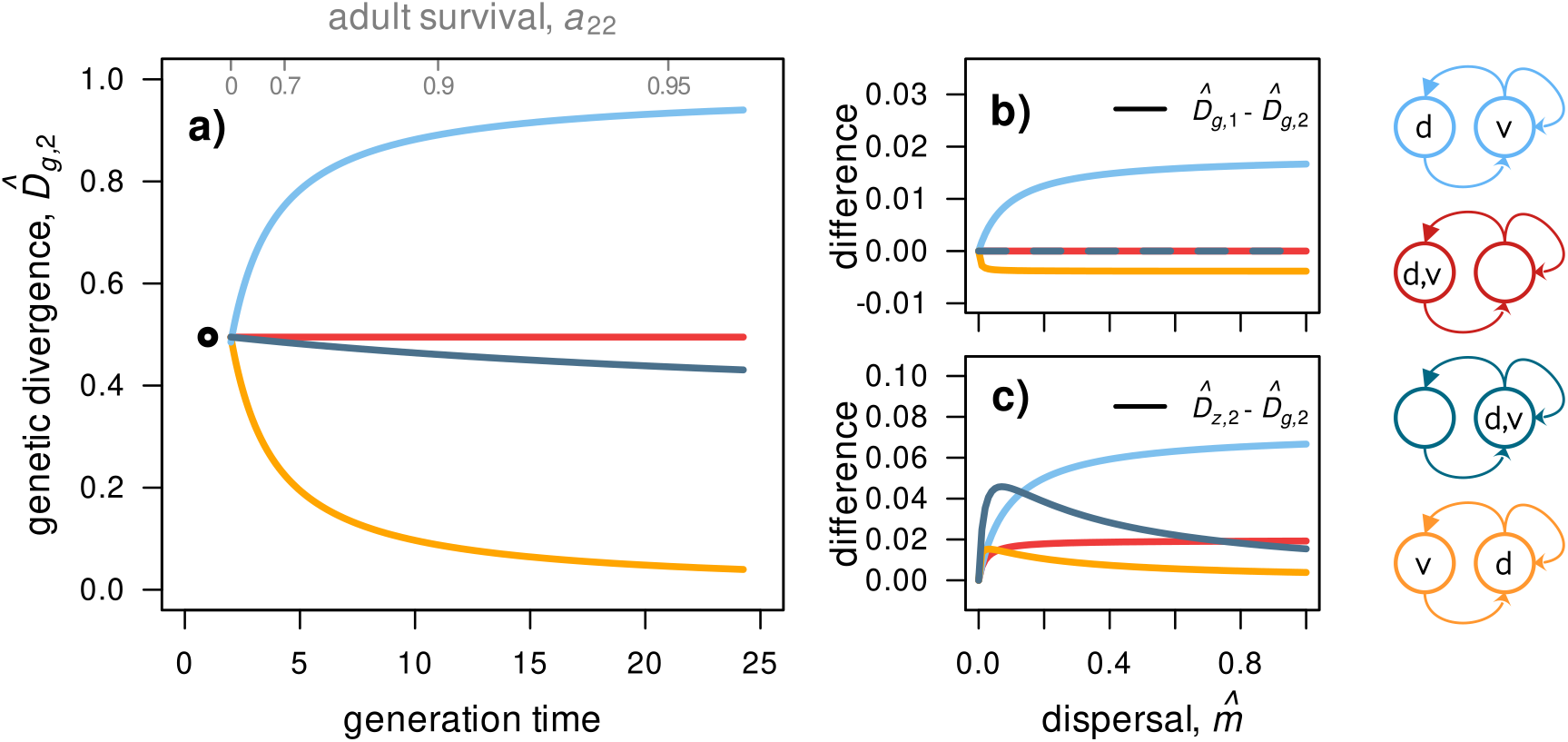
Graph **a)**, genetic divergence in the adult stage (following eqs. 3-6) as a function of generation time (following eq. 2). Graph **b)** illustrates that life stages can differ from each other in the degree of local adaptation when the difference 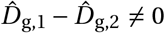. Graph **c)** illustrates a mismatch between genetic and phenotypic divergence with limited heritability (*G*/*P <* 1, see eqs. A15 and A16) such that difference 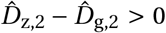. Parameter values: 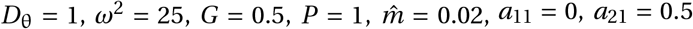. Adult fecundity *a*_12_ is reset for reach parameter combination to achieve *λ* = 1.0. Adult survival in graphs b) and c) is *a*_22_ *=* 0.8. Timings of dispersal and viability selection for each species are indicated with letters *d* and *v*, respectively, in life-history diagrams on the right.

In contrast, when juveniles are exposed to selection but adults disperse (i.e. equilibrium genetic divergence in adults is 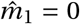 and 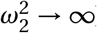),

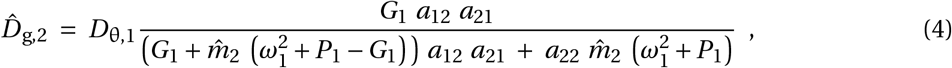

which now depends on vital rates *a*_12_, *a*_21_, and *a*_22_. In particular, as the denominator of eq. (4) increases with adult survival *a*_22_, genetic divergence 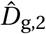 falls below that of an annual species when *a*_22_ *>* 0 (see Fig. 2b, orange line). This is because the lifetime number of dispersal opportunities increases with adult lifespan in this scenario. These repeated dispersal opportunities increase effective dispersal 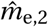 and thus oppose the evolution of genetic divergence. Specifically, 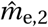 here increases with the per year adult dispersal propensity 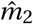 and the number of years spent in that class, which on average is 1/(1 − *a*_22_). With generation time increasing with adult survival *a*_22_ (eq. 2), we find that local adaptation in this case is negatively associated with generation time (see Fig. 3a, orange line). Such a dispersal scenario might apply to species such as amphibians, where adaptation to the local habitat is set by a trait in tadpoles but adults are mobile and move between ponds.

Next, we consider the opposite scenario where it is juveniles that disperse and adults that experience selection (i.e. 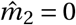 and 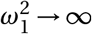).Such a case could apply to trees, where seed dispersal occurs only once, early in life, while grown plants experience viability selection each year anew. Here, equilibrium genetic divergence in adults is

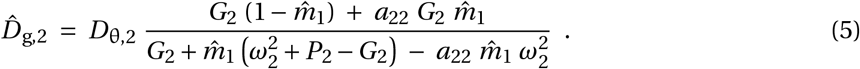

When adults only live for a year (*a*_22_ *=* 0), eq. (5) reduces to eq. (8) in Hendry et al. (2001), which considers an annual species with selection followed by dispersal. When *a*_22_ *>* 0, eq. (5) shows that 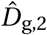 increases with adult survival *a*_22_ (as the denominator gets smaller with *a*_22_). Local adaptation thus increases relative to the annual case with *a*_22_ (see also Fig. 2c, light blue line). This is because adults can survive and reproduce over multiple years in this model, so that they are exposed to viability selection multiple times throughout their lives. Dispersal, in contrast, happens only once, early in life. Selection is therefore reinforced relative to dispersal when *a*_22_ *>* 0 and results in more pronounced genetic divergence in long-lived than annual species. As a consequence, local adaptation is positively associated with generation time here (see Fig. 3a, light blue line).

Finally, we assume that selection and dispersal happen only in adults (i.e. 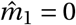 and 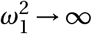).Such a scenario might apply to species such as fish, birds, or mammals, where adults survive over multiple years, experience selection and disperse, while juveniles are sessile and not under selection for the focal trait. Equilibrium genetic divergence in adults is then given by

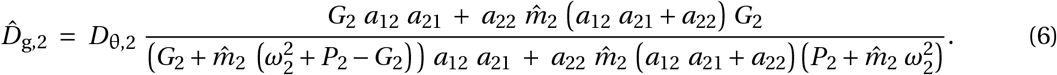

Though this is not immediately obvious from eq. (6), 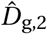 here is only marginally smaller than in the annual case (eq. 3, see Fig. 2d, dark blue line). To understand this, consider that because selection and dispersal opportunities both take place in adults each year, they occur with the same frequency in an individual’s lifetime. As a result, the effects of life-history on effective dispersal and selection cancel each other, bringing genetic divergence close to the annual case. In fact, 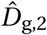 in eq. (6) is equal to eq. (3) under full heritability (*P*_2_ *= G*_2_; also when adults live for a single year *a*_22_ *=* 0). Limited heritability leads to lower genetic divergence because in this case, selection induces a greater phenotypic than genetic response. This greater phenotypic response persists over multiple years when individuals are long-lived (here when *a*_22_ *>* 0), which weakens the genetic divergence and pushes 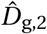 below that of an annual species.

### 3.2 Local adaptation differs between stage classes and does not equate with phenotypic divergence

Our model can also be used to quantify genetic divergence in juveniles 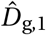 (Appendix A.3.3 for full expressions of 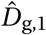). We compare genetic divergence in adults and juveniles in Fig. 3b, which shows the difference 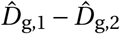 for the life-cycle combinations explored in section 3.1. These comparisons indicate that differences in 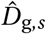 between stage classes evolve although they are typically small. We see the greatest difference under juvenile dispersal and adult selection (i.e. 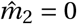 an 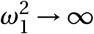), where juveniles evolve larger genetic divergence than adults 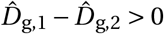, light blue line in Fig. 3b). This is because juveniles experience a different “history” of selection and dispersal than adults here. At census, the adult stage contains many (former) juveniles that dispersed and then matured that same year. Juveniles, in contrast, descend from adults that just experienced divergent selection and then reproduced. This differing history of recent selection and dispersal between juveniles and adults then leads to different levels of genetic divergence at evolutionary equilibrium.

Because phenotypic divergence is much less laborious to measure than genetic divergence in natural populations, it is attractive to use 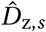 as an alternative measure of local adaptation to 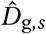. However, comparing genetic with phenotypic divergence under the four different scenarios seen in section 3.1 suggests that 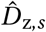 rarely accurately reflects 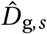. For all four life histories, we find that phenotypic divergence exceeds genetic divergence (i.e.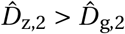) when *G*_*s*_ *< P*_*s*_ (e.g. Fig. 3c). The difference between genetic and phenotypic divergence is especially pronounced when heritability is low (i.e. *G*_*s*_ ≪ *P*_*s*_) and when individuals under selection are long-lived. Under limited heritability, selection elevates phenotypic divergence more so than it increases genetic divergence (see eq. A15 and A16 in Appendix), and this mismatch persists over multiple years when individuals survive. As a consequence, the gap between genetic and phenotypic divergence widens with weak heritability and more frequent iterations of selection on long-lived individuals.

All our results so far assume that juveniles mature in a single year whereas adults are potentially long-lived. We repeated the above analyses for a scenario where it is juveniles that are long-lived (e.g., where plant seeds stay in the seed bank for several years) while adults die after a year (Appendix A, section A.3.4 for analyses). As expected, local adaptation in this case is reduced when selection applies to adults and dispersal occurs at the long-lived juvenile stage. We thus find the same mechanisms being at work in this alternative life-history scenario, even though for a different scheduling of selection and dispersal.

### 3.3 Life-cycle effects on fitness, additive genetic variance, and quantitative trait loci

We also tracked the evolution of local adaptation in individual-based simulations for the different life-cycle scenarios explored in section 3.1 (Fig. 4a, see Appendix B for details). A comparison of 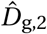 between our mathematical (Fig. 3a) and simulation results (Fig. 4a) shows that these match qualitatively speaking: genetic divergence decreases with generation time when long-lived adults disperse (orange line in Fig. 4a), and increases when selection acts on adults (light blue line in Fig. 4a). Quantitatively however the match between our mathematical and simulation results is weaker. This results from the fact that while additive genetic variance *G*_*s*_ is assumed to be constant in our mathematical model, it evolves in our simulations. This is particularly evident where selection applies to adults, in which case additive genetic variance declines with increasing adult survival as repeated exposure of adults to selection depletes additive genetic variance (Fig. 4c, in blue). As a consequence, when selection and dispersal happen only in adults (dark blue lines in Fig. 3a and Fig. 4a), 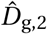 is only weakly affected by adult survival *a*_22_ in our mathematical model (eq. 6, with constant *G*_*s*_), but strongly declines with *a*_22_ in our simulations due to the corresponding decline in *G*_*s*_. Note however that when we plug the values for *G*_*s*_ and *P*_*s*_ that evolve in our simulations into eqs. (3)-(6), mathematical and simulation results match perfectly (Fig. B4 in Appendix B). This confirms that the quantitative mismatch between Figs. 3a and 4a boils down to our assumption that *G*_*s*_ and *P*_*s*_ are fixed parameters in our mathematical approach (as in most evolutionary quantitative genetics).

**Figure 4:**
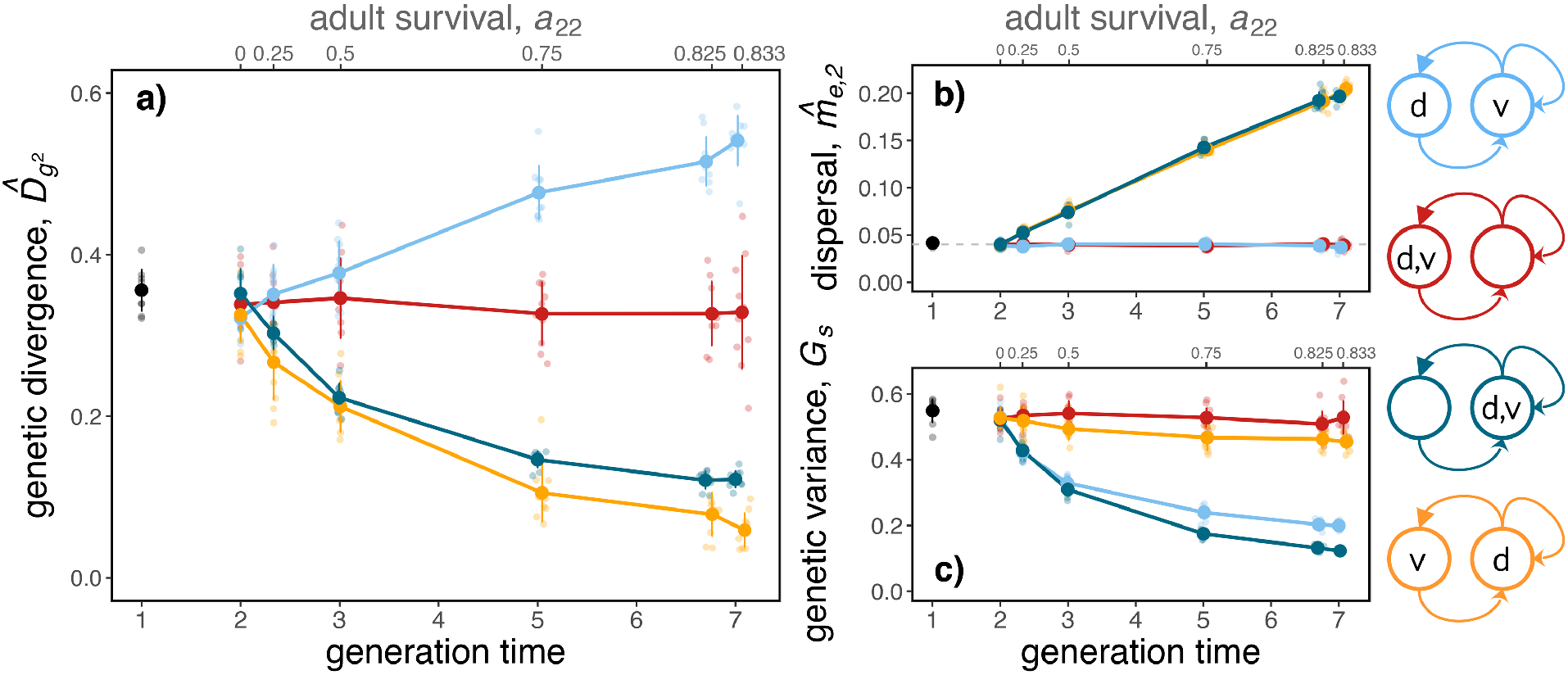
The three graphs show the results of the individual-based simulations. Graph **a)**, genetic divergence in the adult stage 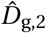 as a function of generation time. Graph **b)**, effective dispersal of adults 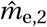 as a function of generation time. The yearly dispersal probability 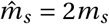 is indicated as a dashed gray line. Effective dispersal 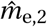 increases with the number of years individuals spend in the dispersing stage (i.e. the stage marked with *d* in the right-hand legend). Graph **c)**, genetic variance *G*_*s*_ before viability selection measured in the stage under selection (i.e. the stage marked with *v* in the right-hand legend). *G*_*s*_ decreases with longevity when adults are under selection. For all graphs: points represent the mean of ten simulations, the error bars correspond to the standard deviations and individual replicates are indicated as small dots. Parameters are: 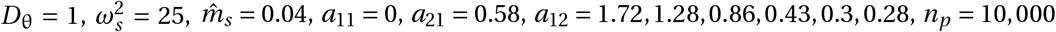.

We also used our simulations to compare the evolved survival *W*_*s,p*_ (*z*_*i,s,p*_) (eq. 1) between local individuals in a habitat and immigrants from the other habitat. As expected, local individuals show greater survival than immigrants across all the different life-cycle scenarios in section 3.1 (Fig. 5a). This is especially the case under scenarios where genetic divergence is large, i.e. when selection applies to long-lived adults and short-lived juveniles disperse (compare light blue box plot in Fig. 5a to light blue line in Fig. 4a). In contrast, we find little fitness differences between locals and potential immigrants when dispersal occurs in adults and these are long-lived, in line with the fact that genetic divergence is low in that case (orange and dark blue box plots in Fig. 5a). Our simulations confirm that this is because effective dispersal 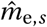 increases with longevity when dispersal occurs each year in adults (orange and dark blue curves in Fig. 4c).

**Figure 5:**
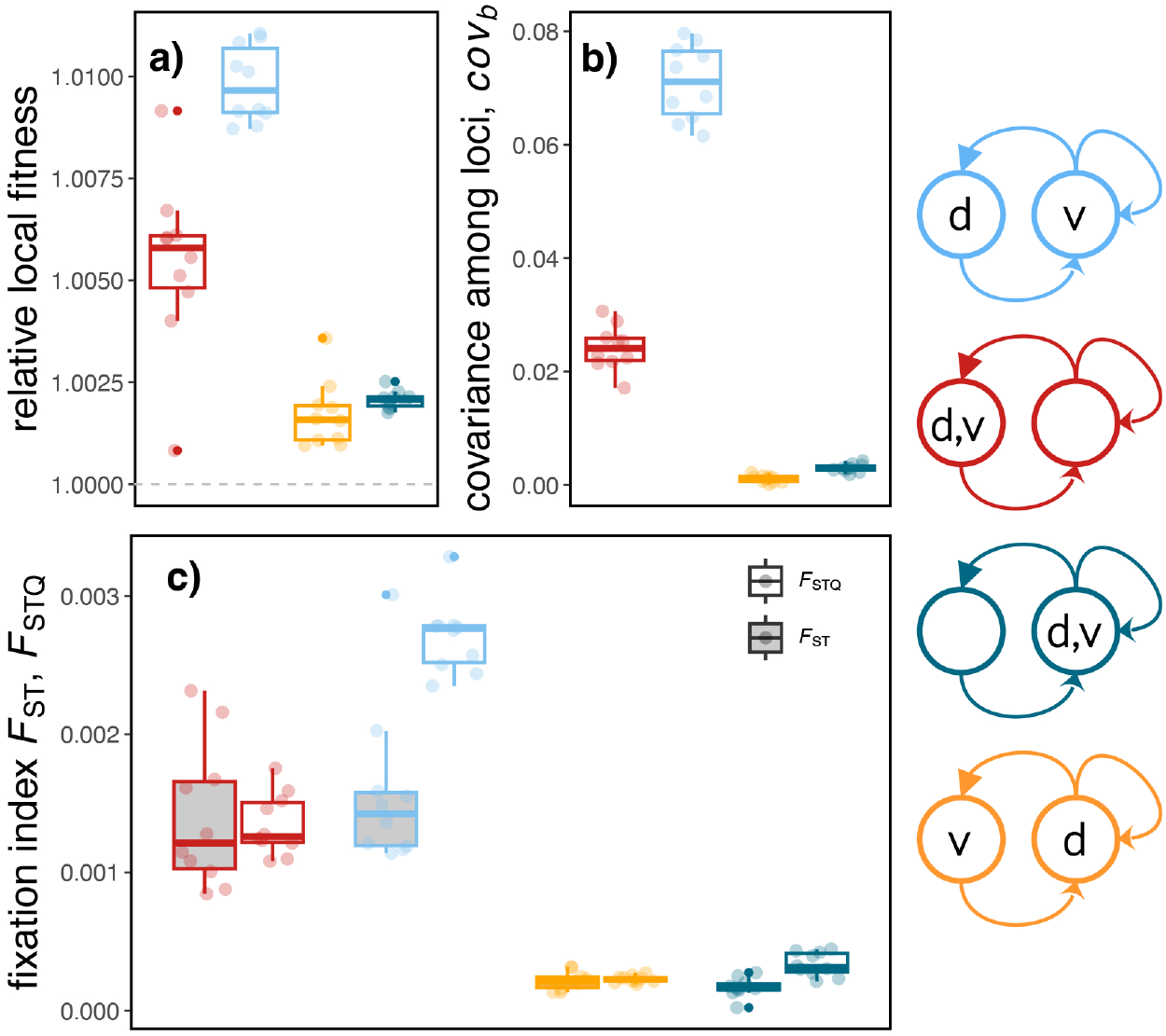
Population statistics collected from the simulations of long-lived species (*a*_22_ *=* 0.833). In graph **a)**, we show the relative fitness of local individuals compared to individuals from the other patch (the absolute survival probability *W*_*s,p*_ (*z*_*i*_) of locals divided by *W*_*s,p*_ (*z*_*i*_) of incoming individuals from the other habitat). A relative local fitness of *>* 1 indicates a greater survival probability of locals as compared to incoming individuals. In graph **b)**, we show the covariance between allelic effects among pairs of loci *cov*_*b*_ between patches. We compute the covariance as 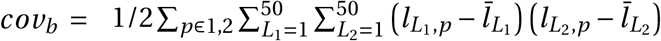 for *L*_1_ ≠ *L*_2_ where 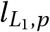 denotes the allelic effect at locus *L*_1_ in patch *p*, and 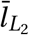 is the allelic effect averaged over both patches. Graph **c)** compares *F*_ST_ for neutral loci to *F*_STQ_ for quantitative loci. Genetic divergence 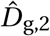 positively correlates with relative local fitness, *cov*_*b*_, and *F*_STQ_ (e.g., compare the light blue life history across plots). The boxplots are generated from ten replicates each. Additional parameters are: 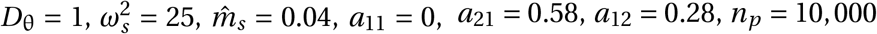. For more on relative local fitness, *cov*_*b*_, and *F*_STQ_, see Fig. B2 and B3 in Appendix B. Timings of dispersal and viability selection for each species are indicated with letters *d* and *v* respectively in life-history diagrams on the right.

Finally, we use our simulations to study how genetic divergence, which reflects divergence at the trait level, associates with genetic responses at single loci. We do so in two complementary ways. First, we measure how allelic composition at single loci differs on average between patches. This can be quantified with the between-patch differentiation in allele frequencies, typically denoted by *F*_STQ_ for quantitative loci and *F*_ST_ for neutral loci (following Le Corre and Kremer, 2012). These are routinely used in empirical studies to investigate the genetic basis of traits (e.g., Hoban et al., 2016). Second, we measure the average covariance *cov*_*b*_ in allelic effects among pairs of loci, which indicates whether trait loci found in the same genomes tend to act in the same phenotypic direction (when *cov*_*b*_ *>* 0; Le Corre and Kremer, 2012). We observe that life-cycle scenarios associated with larger genetic divergence 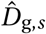 are also associated with larger between-patch differentiation in allele frequency *F*_STQ_ at quantitative trait loci, and with a greater difference between quantitative and neutral loci *F*_STQ_-*F*_ST_ (compare 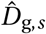 in Fig. 4a to *F*_STQ_ in Fig. 5c; see also Fig. B3). Similarly, these show elevated covariance of allelic effects among pairs of loci *cov*_*b*_. These results suggest that local adaptation under different life histories may leave different signatures at the level of allele frequencies (i.e. *F*_ST_ and *F*_STQ_) and covariance among quantitative trait loci (*cov*_*b*_).

## 4 Discussion

Our analyses indicate that long-lived species with extended generation time readily evolve local adaptation differently than short-lived species (all else being equal). The magnitude and direction of this difference are set by the relative scheduling of viability selection and dispersal. Put simply, local adaptation is enhanced (inhibited) in long-lived species when selection episodes occur more (less) frequently throughout an individual’s life than dispersal opportunities (eqs. 4 and 5). The notion that repeated exposures to selection amplifies evolutionary response compared to selection applied only once (again, all else being equal), can be seen from the general results of Barfield et al. (2011) (their eq. 8; see also Lande, 1982). Here, we have also considered that individuals can disperse between habitats and experience different selection pressures there. As our results reveal, it isn’t just how frequent selection is applied that matters then, but also how frequent it is relative to dispersal. In fact, when the life cycle is such that dispersal and selection balance, e.g. when both selection and dispersal take place in lockstep (eqs. 3 and 6), then local adaptation may be largely insensitive to lifespan.

Two separate meta-analyses of transplant experiments in a wide range of taxa did not detect any significant association between local adaptation and life-history characteristics (e.g., Leimu and Fischer, 2008; Halbritter et al., 2018). This aligns with our results, which support the idea that there should not be a consistent and unidirectional effect of generation time or iteroparity across taxa on genetic divergence between habitats. Instead, long-lived iteroparous species may well evolve greater, weaker, or even similar levels of local adaptation than short-lived semelparous species, depending on the context of selection and dispersal. More generally, it is good to remind ourselves that local adaptation is determined by many different factors (such as gene flow, genetic correlations among traits, or pheno-typic plasticity; e.g., Hendry et al., 2001; Guillaume, 2011; Schmid and Guillaume, 2017) and that all these factors vary between species and study sites. This poses considerable challenges in detecting statistical associations between local adaptation and life-history characteristics across study systems.

Local adaptation is positively associated with generation time when greater longevity involves more episodes of selection relative to dispersal (Fig. 3a or eq. 5). This result may help understand why trees, which typically have long generation times and strong gene flow (owing to extensive pollen dispersal), nevertheless often show pronounced local adaptation (Howe et al., 2003; Petit and Hampe, 2006; Savolainen et al., 2007). Empirical studies have typically attributed this to strong selection (e.g., Petit and Hampe, 2006). Our findings support this view and identify two routes to strong selection. First, strong selection can simply come from individuals being exposed to strong selection once in their lifetime (i.e. large mortality due to maladaptation, Hendry et al., 2001). Second, strong selection can arise from moderate and even weak selection each year, provided an individual is exposed to such selection repeatedly (as shown in Fig. 2c). In trees, for instance, local adaptation is often set by cold hardiness, which determines individual survival each winter and early spring (Howe et al., 2003). Thus, even weak selection within a year can leave signatures of local adaptation at the gene and fitness level when iterated multiple times.

In contrast, local adaptation and generation time are negatively associated when greater longevity involves fewer episodes of selection relative to dispersal (Fig. 3a or eq. 4). Here, a species’ life history elevates gene flow beyond annual dispersal when individuals can disperse over multiple years. That dispersal can be decoupled from actual gene flow is well-established, though it is typically invoked to explain decreased gene flow between distinct habitats thus favouring local adaptation. Some of the mechanisms that have been proposed to decouple dispersal and gene flow are: where dispersers experience reduced survival and fecundity in novel environments (leading to isolation by adaptation, Nosil et al., 2008); where local individuals mate preferentially with locally adapted phenotypes, due to e.g. assortative mating based on a so-called magic trait (Servedio et al., 2011); or where dispersal is directed towards similar environmental conditions but diminishes along environmental clines (i.e. matching habitat choice; Edelaar et al., 2008). Our results add another, perhaps more straightforward, mechanism to this list: where gene flow is elevated beyond annual dispersal propensity due to repeated dispersal opportunities in long-lived species (Fig. 4b). When long-lived individuals can disperse each year anew, genetic exchange between habitats is multiplied, thus boosting gene flow among sites.

Our model also helps refine the connection between genetic and phenotypic divergence among different habitats. Local adaptation in our model, that is genetic divergence between patches, does not fully coincide with phenotypic divergence (Fig. 3c or Fig. B5). It is well known that phenotypic divergence is an insufficient predictor of local adaptation, but this mismatch is commonly assigned to the effect of phenotypic plasticity (e.g., with counter-gradient variation, Conover and Present, 1990; Conover et al., 2009; Schmid and Guillaume, 2017). In our model, where there is no phenotypic plasticity, the mismatch between genetic and phenotypic divergence unfolds from life history. Where heritability is low, viability selection leads to greater responses at the phenotypic level as compared to the genetic level (e.g., compare eq. A15 to eq. A16 in Appendix A). But whereas this mismatch is reset after reproduction (when the average phenotype in juveniles is reset to their average genetic value), it persists and is amplified when individuals survive over multiple years. This is similar to the mechanism described in Orive et al. (2023) where clonal offspring partially inherit the environmental trait component *e* from their parent. Beneficial environmental effects can thus persist for several generations, and lead to greater phenotypic than genetic divergence, even in absence of phenotypic plasticity.

Our results suggest that elaborate and labor-intensive study design may be unavoidable to study local adaptation in long-lived species. The meta-analyses of Fraser et al. (2011), Halbritter et al. (2018), and Runquist et al. (2020) report that currently available empirical data on long-lived species primarily concern early stage classes (seedlings or juvenile stages). This should limit our ability to detect local adaptation. For instance, where survival differences within a year are small, short-term data (e.g. over one year) would fail to detect the build up of local adaptation through repeated exposures to selection (as seen in see Fig. 5). In other words, because it can take many years before local adaptation becomes detectable at the fitness level (for a telling example, see Germino et al., 2019), it would be useful if data over multiple stages – ideally over the whole lifespan of individuals – were available (Germino et al., 2019; Wadgymar et al., 2022). Having such data would also help us to quantify the causal factors of local adaptation, that is selection and dispersal. There are at least two ways to go about this. First, one could estimate selection and dispersal over the course of an individual’s lifespan, which would directly account for potentially cumulative effects of repeated selection and dispersal. However, obtaining such longitudinal data that combines information on selection and dispersal over multiple years remains a challenging endeavour (Svensson, 2023). Alternatively, and probably more easily, one could collect data on dispersal and selection for multiple individuals that belong to multiple stages over a single year, for instance from cross-sectional data (e.g., Svensson, 2023). These data could then be integrated like in our model, which uses annual selection strength and annual dispersal probabilities as input, to better understand local adaptation in long-lived species.

Several other processes and mechanisms that were ignored in our model may further complicate the relationship between life history and local adaptation. Ontogenetic trait changes (Saltini et al., 2023) or stage-specific plastic responses (Fischer et al., 2014), for instance, might either enhance or inhibit local adaptation depending on the context in which they occur. Additionally, if local adaptation depends on multiple correlated traits, these traits could co-vary genetically within and/or between stages and thus influence the evolutionary response (Guillaume, 2011; Barfield et al., 2011). For small populations, it has been suggested that long-lived life histories can influence local adaptation through their effects on the effective population size (Leimu and Fischer, 2008). Age- or stage-structured life histories can show a reduced effective population size compared to short-lived species depending on life-cycle events, and thus limit the efficacy of selection in certain cases (Charlesworth, 2009). Therefore, considering these aspects should further reinforce the message of our results: that a species’ life history may not have a simple, unidirectional effect on the evolution of local adaptation.

## 5 Author contributions

L.H., C.M., and M.S. conceived the project. M.S. performed the mathematical analyses with contributions from L.H. and C.M., and L.H. ran the simulations with input from M.S. and C.M. All authors contributed to manuscript writing.

## 6 Acknowledgments

C.M. is funded by Swiss National Science Foundation grant PCEFP3181243.

## 7 Conflict of interest

The authors declare no conflict of interest.

## 8 Data and code availability

The simulation code and data will be made publicly available upon acceptance on Figshare.

## A Analytical model

### A.1 Trait divergence and local adaptation

We study the evolution of local adaptation at the example of a single species in a two-patch setting. The focal species has a stage-structured life history with a total of *k* stage classes. Local adaptation in a stage indexed *s* is found when the species adapts genetically to the local environment and evolves genetic divergence 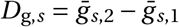 and phenotypic divergence 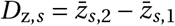 between patches (see section 2.2 in the main text for more details on these metrics). We collect all values of *D*_g,*s*_ and *D*_z,*s*_ in the column vector

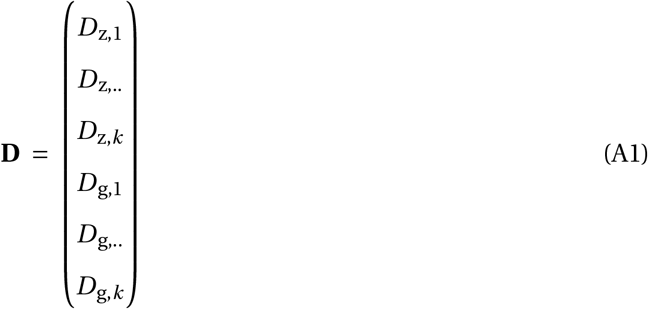

with 2*k* entries. The first *k* entries of **D** collect phenotypic divergences at all stage classes, the last *k* entries contain genetic divergence.

Within each year, genetic and phenotypic divergence between both patches change from three events: dispersal (d), viability selection (v), and life-history transition (t). At evolutionary equilibrium, the vector of genetic and phenotypic divergence **D** is constant among years when the consecutive effects of life-history transition, migration, and selection balance out (we denote **D** at migration-selection-transition balance as 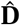). To compute 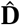, we first derive the recursion equation

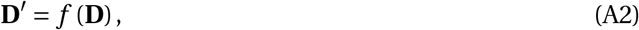

to compute phenotypic and genetic divergence in year *τ +* 1 (**D**^*′*^) as a function of divergence in the previous year *τ* (**D**). In a second step, we solve eq. (A2) at equilibrium, when 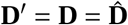. The order of events matters, and we account for all six combinations of dispersal, selection, and life-history transitions (tdv, tvd, dtv, dvt, vtd, vdt). For instance, we study the dvt sequence of yearly events when dispersal is followed by viability selection and then life-history transition.

### A.2 Analysis

In the following sections, we specify how each yearly event (dispersal, viability selection, and life-history transitions) modifies the genetic and phenotypic divergence between patches.

#### A.2.1 Dispersal

We assume that individuals of stage *s* disperse to the other patch with an annual dispersal probability *m*_*s*_ and stay philopatric with probability 1 − *m*_*s*_. The annual dispersal probabilities are symmetric, such that the probability of stage *s* individuals to leave patch 1 is identical to the probability of leaving patch 2. The average phenotypic and genetic values in both patches before dispersal 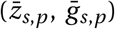 change with dispersal to

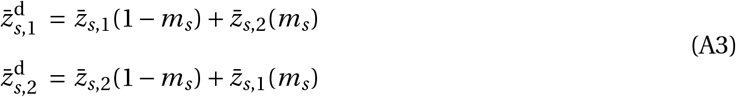

and

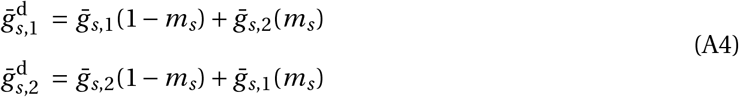

where the superscript *d* denotes metrics after dispersal. From eq. (A3) and (A4), we derive the phenotypic and genetic divergence after 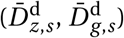, as

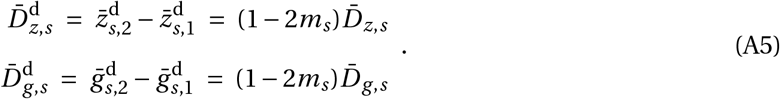

Equations (A5) can be expressed in matrix form as

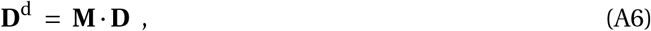

where the vector **D** immediately before dispersal is left multiplied by the 2*k*-by-2*k* matrix

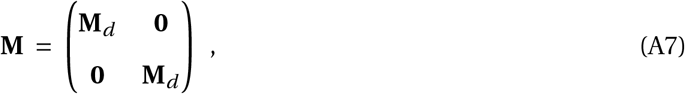

with

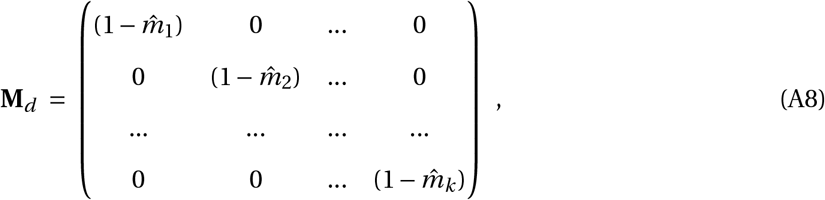

where 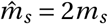. More formally, we express eq. (A6) as a function

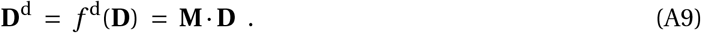

In the absence of individual dispersal (when all 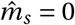), matrix **M** reduces to the identity matrix and the entries of **D** do not change during this yearly event (**D** *=* **D**^d^). In a panmictic population, when half of the individuals disperse to the other patch (where all *m*_*s*_ *=* 0.5 and thus 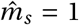), matrix **M** only contains zeros and dispersal eliminates phenotypic and additive genetic divergence altogether. Intermediate dispersal reduces genetic and phenotypic divergence to some extent.

#### A.2.2 Viability selection

Each stage can be subject to viability selection. Individual *i* in stage *s* and patch *p* survives viability selection with probability

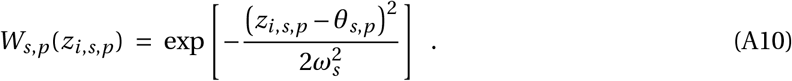

Applying viability selection to all individuals of a certain stage class might lead to a change in the average phenotype (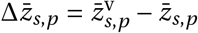, where superscript v denotes metrics after viability selection). The change in the average phenotype from viability selection was derived by Lande and Arnold (1983) as

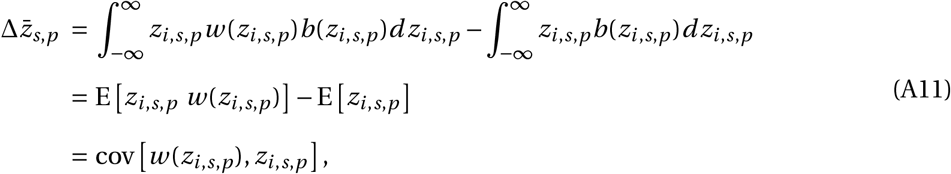

where the average phenotype before and after selection is computed from the product of the individual trait value (*z*_*i,s,p*_), relative survival (*w* (*z*_*i,s,p*_)), and the frequency of z-values (*b*(*z*_*i,s,p*_)), integrated over the phenotypic distribution. Relative survival 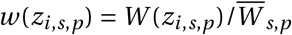 denotes the fraction of the phenotypes absolute survival probability *W* (*z*_*i,s,p*_) (following eq. A10) over the average survival within stage *s* and patch 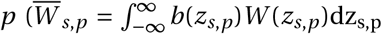,where *b*(*z*_*s,p*_) denotes the frequency of phenotype *z*_*s,p*_). Eventually, the phenotypic change in patch *p* and stage *s* from viability selection corresponds to the covariance between the individual trait values *z*_*i,s,p*_ and their relative survival probabilities *w* (*z*_*i,s,p*_).

When assuming that phenotypic and additive genetic values within each patch and stage follow a bivariate Normal distribution, a linear relationship between the individual *z*- and *g*-values follows. This linear relationship has a slope of *G*_*s,p*_ /*P*_*s,p*_, the fraction of the additive genetic variance over the phenotypic variance (i.e. the heritability of the trait). As a consequence, selection on the phenotype also causes a change in the average additive genetic effect

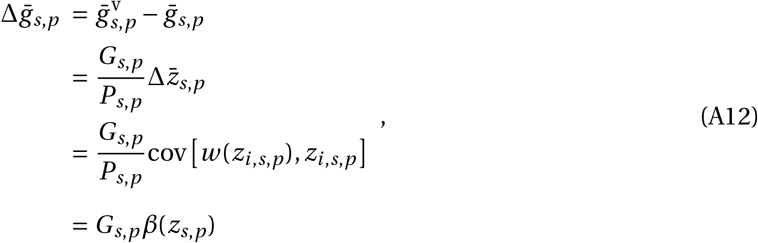

where *β*(*z*_*s,p*_) *=* cov *w* (*z*_*i,s,p*_), *z*_*i,s,p*_ /var *z*_*s,p*_ is the partial regression coefficient of relative survival on the trait value. Lande (1979) and Via and Lande (1985) derived the partial regression coefficient as

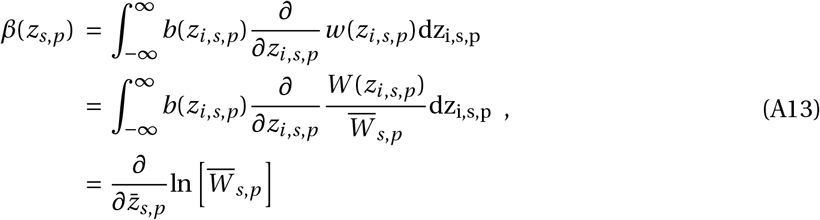

where 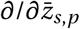 is the gradient operator with respect to the trait mean evaluated at the trait mean. For normally distributed phenotypic and genetic values with a Gaussian survival function, eq. (A13) becomes

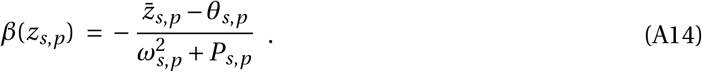

From eqs. (A11)-(A14), the phenotypic change from one round of viability selection becomes

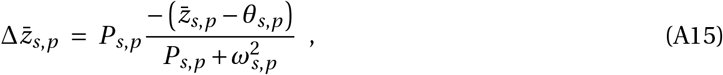

and the change in the average breeding value becomes

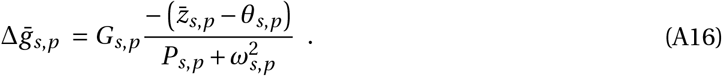

From eq. (A15) and (A16), we then express the change in phenotypic and genetic divergence between patch *p =* 1 and *p =* 2 from viability selection as

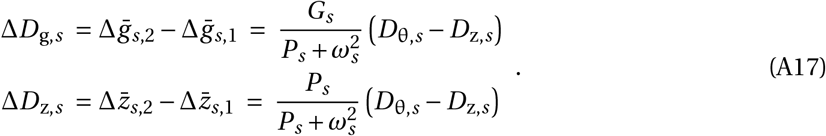

Here, we assume that *G*_*s*_ and *P*_*s*_ are identical in both patches and that the optimal phenotypic divergence for each stage 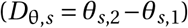 is centered around that same average value 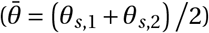.

Equation (A17) can be expressed in matrix notation as

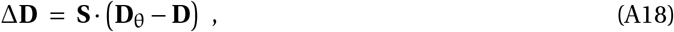

with the 2*k*-by-2*k* matrix

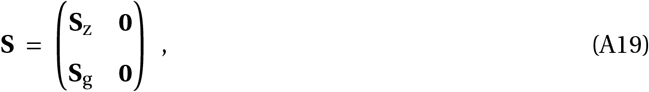

and the *k*-by-*k* matrices

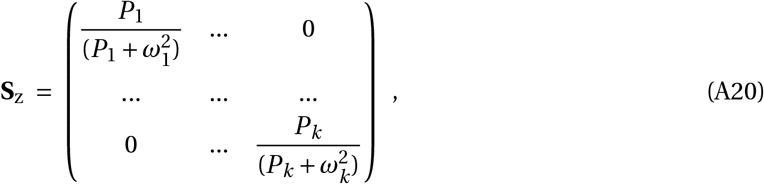

and

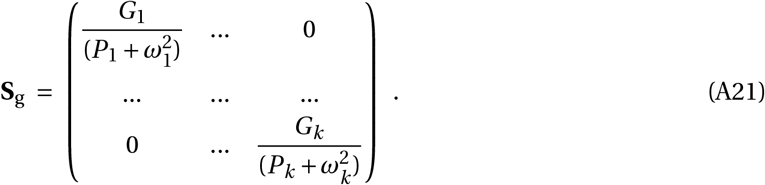

The column vector

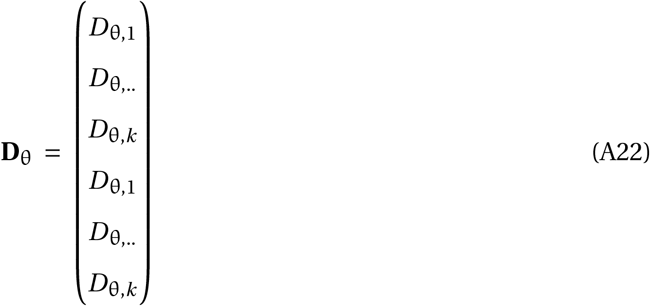

holds the stage specific values of optimal divergence. Following from eq. (A18) we arrive at function

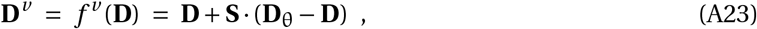

where genetic and phenotypic divergence immediately before viability selection **D**, together with vector **D**_θ_ and the matrix **S**, allow to compute the divergence vector after viability selection **D**^*v*^.

#### A.2.3 Life-history transitions

##### Divergence after stage transitions

From conception to death, individuals pass through a series of stage classes. When stage classes differ from each other in the average genetic and phenotypic composition, transitions from one stage to the other could lead to phenotypic and genetic changes within patches, and modify 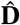.

The average additive genetic value in stage *s* and patch *p* after one round of life-history transition is

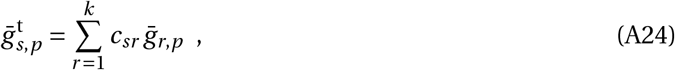

the sum of average additive genetic values in all stages indexed *r* before life-history transition 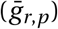, weighted by the relative contribution *c*_*sr*_ of stage *r* individuals to stage *s* (where 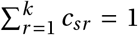). We outline the derivation of *c*_*sr*_ later in this section.

The average phenotypic value of stage *s* individuals after life-history transition is

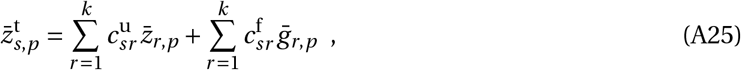

where the first term on the right-hand side of eq. (A25) captures *direct transitions* and the second term reflects *reproductive transitions*. Direct transitions occur when individuals survive the event “life-history transition” to the next year, and then either stay in the same stage or transit to another stage. Reproductive transitions happen when stage *r* individuals contribute offspring to stage *s* by sexual reproduction. The distinction between direct and reproductive transition is necessary because the average phenotype of stage *s* individuals results from the average *phenotypic* value of all contributing stages with direct transitions (the first part of eq. A25), but attributes to the average *additive genetic value* of the parental stage *r* with reproductive transitions (recall that phenotype expression at birth is *z = g +e* with the average environmental effect being zero). Eqs. (A24) and (A25) further indicate that we assume no ontogenetic trait changes. To distinguish between direct and reproductive transitions, we decompose the total relative contributions *c*_*sr*_ (as used in eq. A24) into a component of direct transition 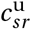 and in one of reproductive transitions 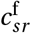 where 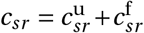. We present the derivation of 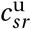 and 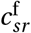 later in this section.

From eq. (A24) and (A25), we obtain how genetic and phenotypic divergence between patches changes from life-history transition

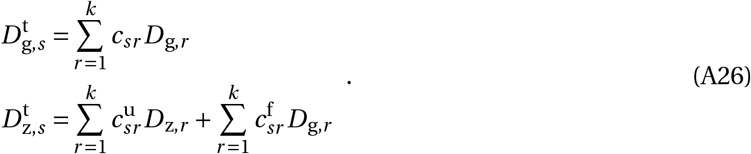

To express eq. (A26) in matrix form, we collect all values of 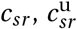, and 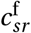 in *k*-by-*k* matrices **C, C**_u_, and **C**_f_, respectively, and combine them into the 2*k*-by-2*k* matrix

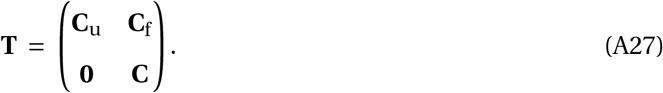

Matrix **T** computes the change in the average phenotypic and genetic divergence from direct and reproductive transitions (**D**^t^ *=* **T D**) and gives rise to the function

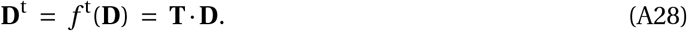

Note that **T** is identical to matrix **K** in Barfield et al. (2011) (their eq. B1).

##### Relative stage contributions

The values 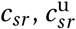 and 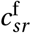 are key to compute the effect of life-history transitions on local adaptation, and we now outline their computation. To this end, we assume that the matrix population model within each patch at evolutionary equilibrium is

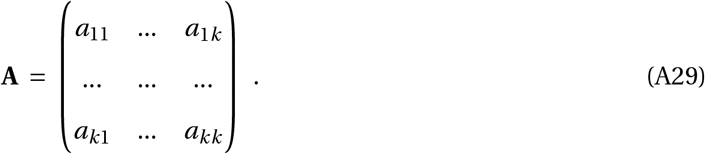

The entries of **A** do not vary with the trait value *z*, and are constant over time. We further assume that matrix **A** and the vector of stage densities **n** *=* (*n*_1_, …, *n*_*k*_)^*T*^ are the same in both patches (see section 2.1 of the main text), and that the species in each local patch reached its stable stage distribution (SSD) as captured by **q**, a column vector containing the frequency of stage *s* individuals *q*_*s*_ (with *q*_*s*_ *=* 1). Recall that dispersal is symmetric between patches and selection is weak such that these processes do not distort the entries of **q**. Each patch further reached the asymptotic growth rate *λ*, the leading eigenvalue of **A**. Most realistically, the growth rate at migration-selection-transition balance is close to *λ =* 1, even though we provide a general solution for *λ* ≥ 1. Importantly, each stage eventually grows at the same rate 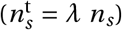. Then, the change in the density of stage *s* from a single event of life-history transition is 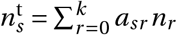 and can be reformulated as

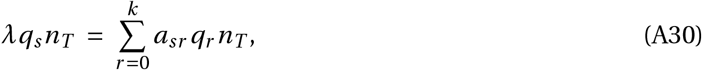

with the total population size *n*_*T*_ *= n*_1_ *+n*_2_ *+*…*+n*_*k*_, and the number of stage *r* individuals being *q*_*r*_ *n*_*T*_. From eq. (A30), we obtain the relative contribution of stage *r* individuals to stage *s* as

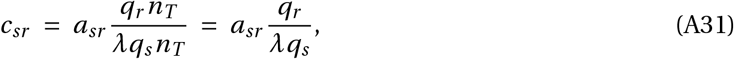

and collect these values in matrix **C**

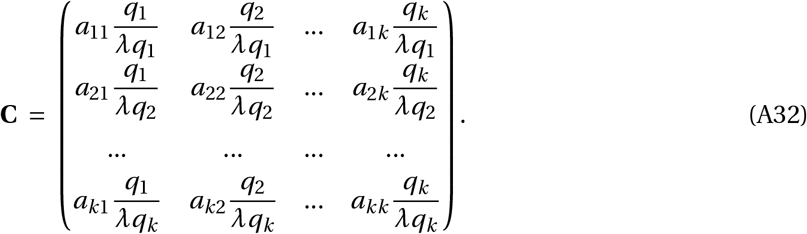

Note that matrix **C** is identical to matrix **C** in Barfield et al. (2011).

To derive matrices **C**_u_ and **C**_f_ as part of eq. (A27), we decompose the entries of matrix **A** into the contributions from direct transitions and reproductive transitions as *a*_*sr*_ *= u*_*sr*_ *+ f*_*sr*_, and use these terms to compute the entries of **C**_u_ and **C**_f_ as

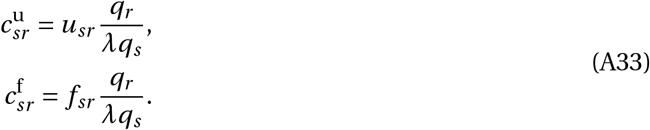

The exact partitioning of **C** into **C**_u_ and **C**_f_, however, is specific to each species and cannot be generalized at this point.

#### A.2.4 Trait divergence at migration-selection-transition balance

After specifying the effect of dispersal, viability selection, and life-history transitions on genetic and phenotypic divergence, we now derive the recursion **D**^*′*^ (eq. A2). We first outline the approach at the example of the dvt life-cycle combination, when each year dispersal occurs before viability selection and life-history transitions.

By help of the functions (A9), (A23), and (A28) we obtain the recursion

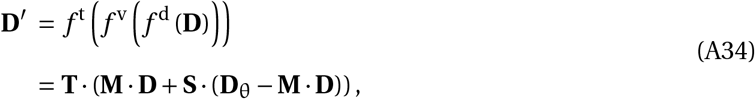

which we re-formulate as an affine model

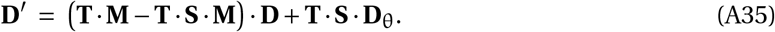

and solve at equilibrium for 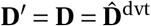. The resulting column vector

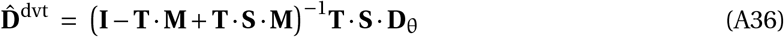

contains the stage-specific genetic and phenotypic divergence between both patches when dispersal is followed by selection and then life-history transitions. Beside the already mentioned matrices **T, S** and **M**, equation (A36) is a function of the identity matrix **I** and the inverse of matrix **I** − **T** · **M** *+* **T** · **S** · **M** (denoted by ^−1^). The analytic solution (A36) is valid for any life-history strategy, with any number of life stages, that can be described by a matrix population model (i.e. a Leslie or Lefkovitch matrix **A**, see section A.2.3), and for which a solution to (**I** − **T** · **M** *+* **T** · **S** · **M**)^−1^ can be found.

Following the same procedure as applied in eqs. (A34)-(A36), we obtain the equilibrium vector of divergence for the other life-cycle combinations as:

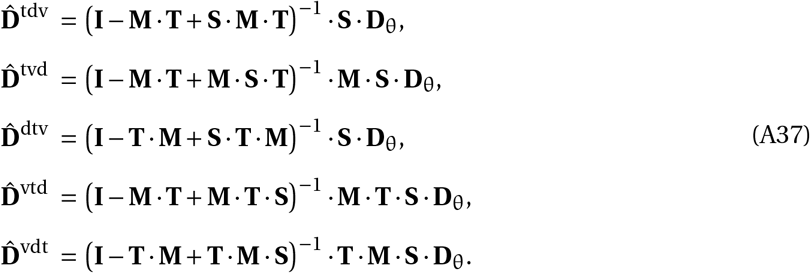

The species’ life history shapes the evolution of local adaptation through the yearly event “*life-history transition*” when the entries of the matrix population model **A** (see section A.2.3) shape **T** and eventually 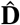. In addition, dispersal and viability selection can act in a stage-specific manner, where **S** and **M** set the intensity and timing of dispersal and viability selection throughout an individual’s life. The resulting vector 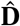 thus is a function of the species’ vital rates *a*_*sr*_, the resulting stable stage distribution *q*_*r*_, and the asymptotic growth rate *λ*. As a consequence, the results depend on life history properties similar to classic findings in life-history theory where trait changes directly link to *a*_*sr*_, *q*_*r*_, and *λ* (e.g., eq. 8 in Barfield et al., 2011). However, we do not model life-history evolution per se but study the evolution of local adaptation while keeping life history constant (i.e. with the entries of **A** being parameter in our models).

### A.3 Results

#### A.3.1 Genetic and phenotypic divergence in an annual species

To illustrate the effect of a species’ life history on the evolution of local adaptation, we first consider a species with a single stage class and non-overlapping generations. All individuals reach adulthood, produce juveniles by sexual reproduction, and die over the course of a single year. Such a life history could refer to an annual plant without seed dormancy, or a univoltine insect. We focus on a scenario where dispersal is followed by selection and life-history transitions within each year (the dvt scenario).

##### Life-history parameters

With a single life stage and non-overlapping generations, the matrix population model is

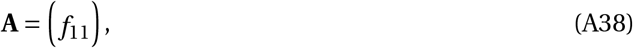

a 1-by-1 matrix where *f*_11_ is the average adult fecundity. This entry *f*_11_ also corresponds to the population growth rate

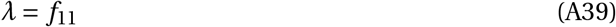

and leads to the stable-stage structure

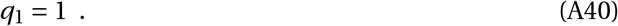

As a consequence, the single entry of matrix **C**_u_ (that contains only direct transitions, see by eq. A27) is zero because no individual survives to the next year. Instead, the entry of matrix **C**_f_ is 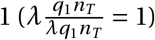 such that **C** *=* **C**_f_ and the matrix **T** (eq. A27) becomes

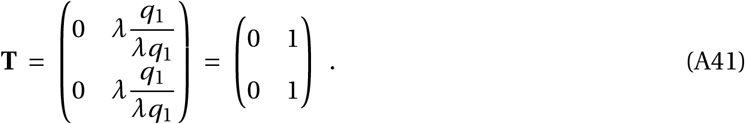

With a single life stage and non-overlapping generations, matrix **M** (A7) becomes

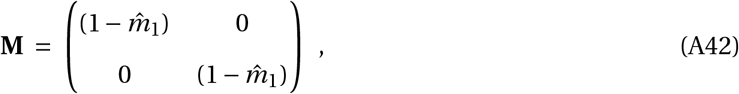

and matrix **S** (A19) is

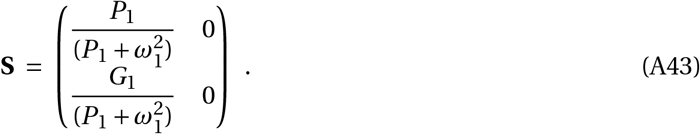

##### Local adaptation

When putting eqs. (A41)-(A43) into eq. (A36), we recover eq. (3) of the main text

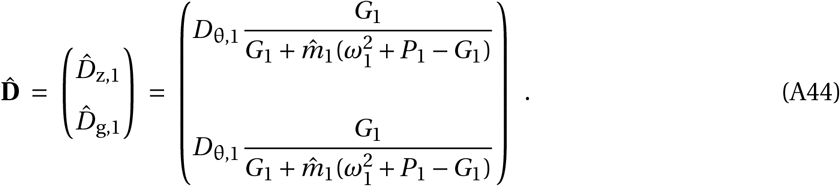

Note that 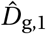 as part of vector (A44) is identical to eq. (7) of Hendry et al. (2001) who derived genetic divergence for an annual species when dispersal is followed by selection.

When we instead assume a different order of yearly events, and plug eqs. (A41)-(A43) into eq. (A37) for the vtd life-cycle combination, we recover

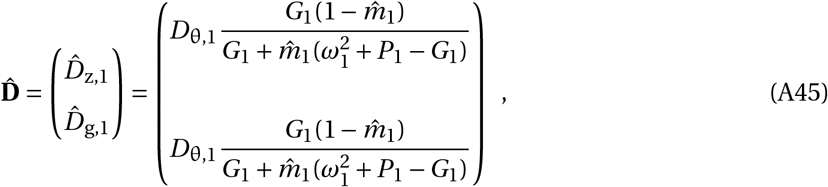

which is identical to eq. (8) of Hendry et al. (2001).

#### A.3.2 Local adaptation with a two-stage life history

##### Life-history parameters

As one example of a long-lived species, we consider a species with two life stages, where individuals could either be in the juvenile (*s =* 1) or in the adult stage (*s =* 2). During *life-history transition*, juveniles stay in the juvenile stage with probability *u*_11_ and transit to the adult stage with probability *u*_21_ (where 0 *<* (*u*_11_ *+ u*_21_) ≤ 1). Each adult individual produces on average *f*_12_ juveniles by sexual reproduction, and each adult survives to the next year with probability *u*_22_. Such a 2-stage life history can be represented by matrix population model

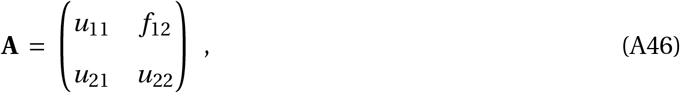

with stage proportions

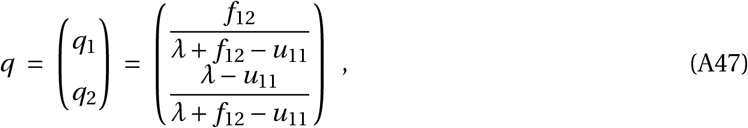

and with an asymptotic population growth rate

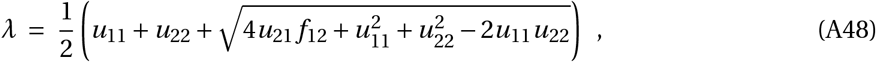

Next, we compute the generation time *T* for a species with life history **A** (eq. A46) that reached its stable stage structure (eq. A47) and is at carrying capacity (when *λ =* 1). We refer to generation time as the average age of parents siring new offspring, and compute *T* from eq. 5.77 in Caswell (2001). This equation applies to age-structured life histories, but can be adapted for matrix **A** (eq. A46) as

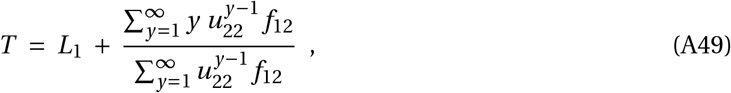

with the average age of juveniles entering the adult stage

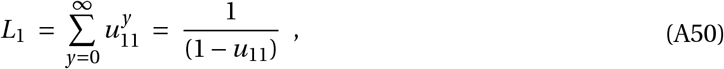

and the (possible) number of years spend as a reproducing adult *y*. Since adult fecundity *f*_12_ is a parameter that does not change with age *y*, eq. (A49) can be simplified to

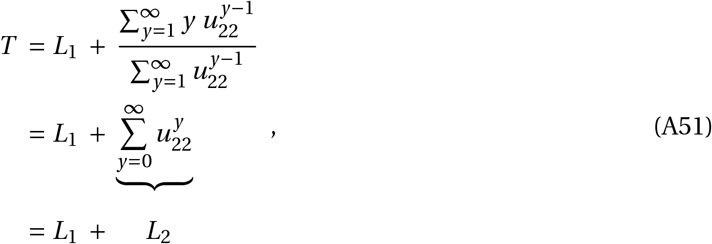

with the expected lifespan as an adult *L*_2_. The generation time thus reduces to

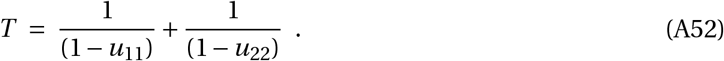

Recall that the vital rates *u*_11_ and *u*_22_ refer to *direct transitions* and here are identical to the entries *a*_11_ and *a*_22_ of matrix **A**, respectively.

The effect of life-history transitions on the divergence vector

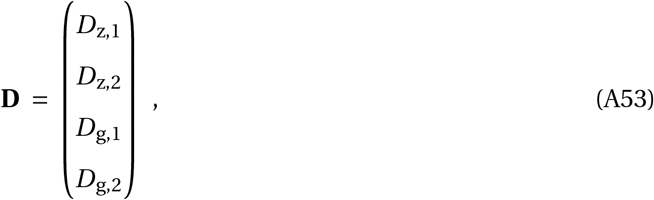

then is captured by matrix

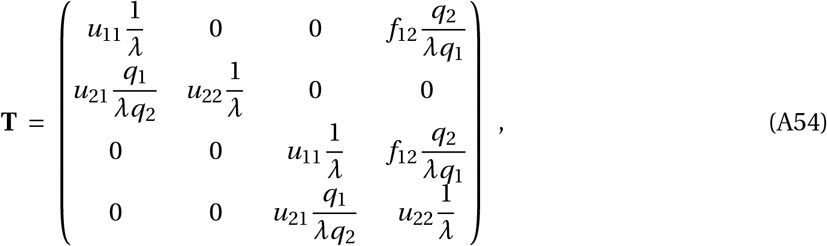

while stage-specific migration is realized by

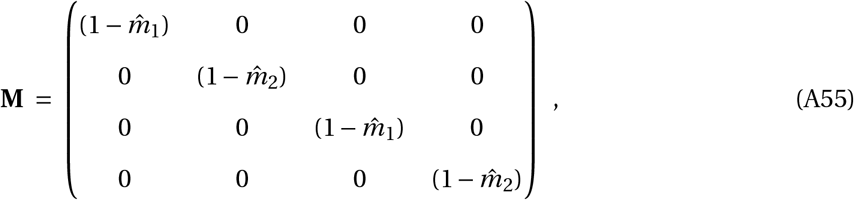

and stage-specific selection links to

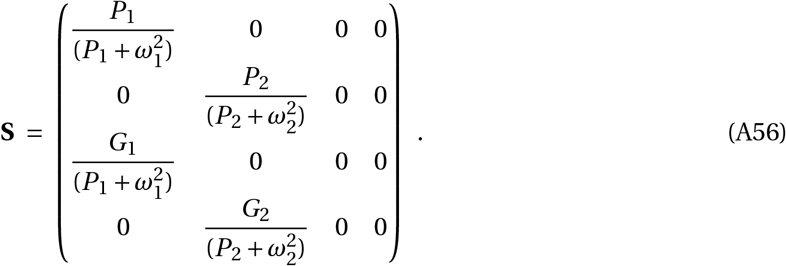

Putting eqs. (A54)-(A56) into eq. (A36) or eq. (A37), we obtain the vector of phenotypic and genetic divergence at migration-selection-transition balance. While this approach allows to compute the extent of local adaptation for a set of given parameter values, the general solution of this vector is rather unsightly, and does not reveal easily how life-history characteristics link to the evolution of local adaptation.

#### A.3.3 Long-lived adults

To reveal the basic underlying mechanisms, we focus on a species with iteroparous long-lived adults (*u*_22_ ≥ 0) but short-lived juveniles (*u*_11_ *=* 0). For this life history, we study how the timing of dispersal and selection affect the evolution of genetic divergence and local adaptation. If one of the two stages is not dispersing (for instance the juvenile stage), we set the migration probability to zero (for instance 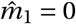). When the trait is not under selection in a stage class (for instance in the adult stage), and the selection gradient becomes zero, we set the genetic and phenotypic variance to zero (e.g., *G*_2_ *=* 0 and *P*_2_ *=* 0), which is mathematically equivalent to an infinitely broad fitness function with 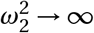 (see eq. A15 and eq. A16).

##### Dispersal and selection only in juveniles

When selection and migration happens only in the juvenile stage (i.e. 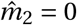 and 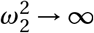),we find that genetic and phenotypic divergence becomes

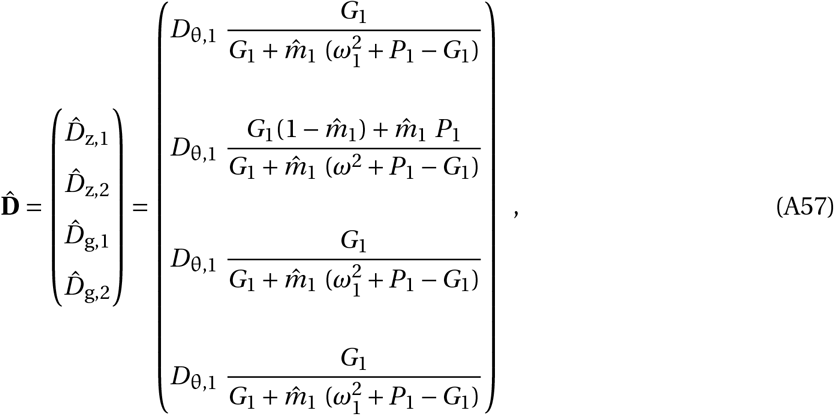

and recover eq. (3) of the main text.

##### Juveniles under selection and adults disperse

With adult dispersal and juvenile exposure to selection (i.e. 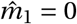 and 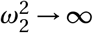),while juveniles are sessile and adults are not under selection, genetic and phenotypic divergence becomes

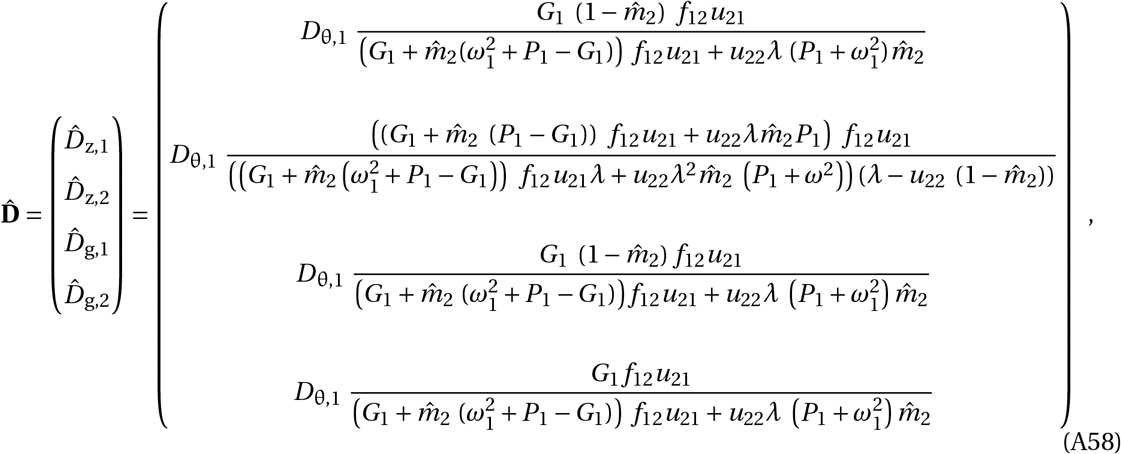

and we thus derive eq. (4) of the main text.

##### Juvenile dispersal and adult selection

When selection acts only on adults, while only juveniles disperse (i.e. 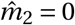 and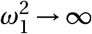), we obtain

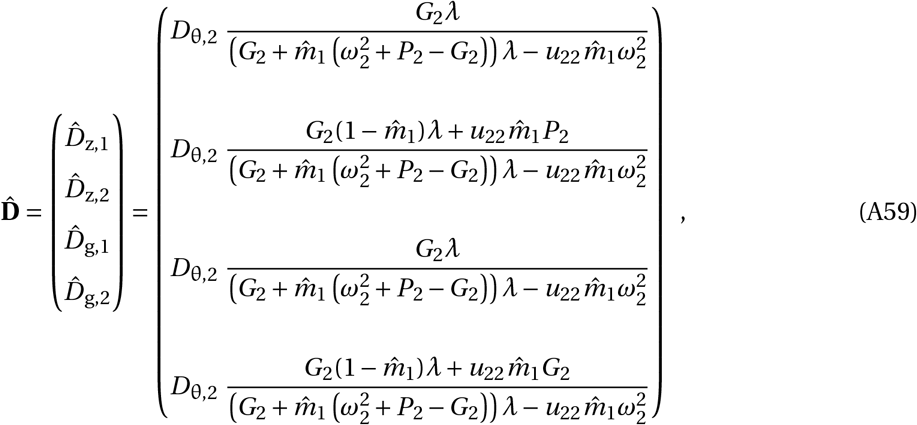

and get eq. (5) of the main text.

##### Adult dispersal and adult selection

When selection and dispersal acts only on adults (i.e. 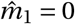 and 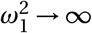), we obtain

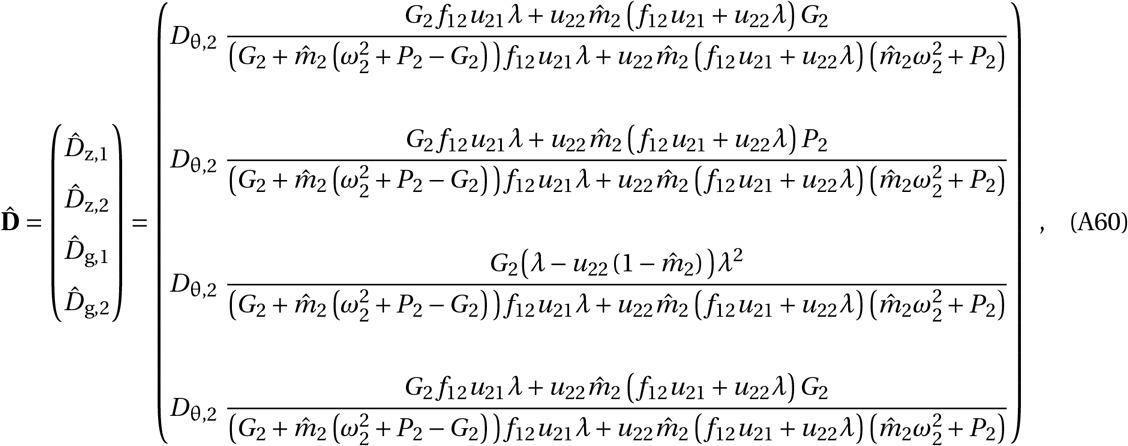

and retrieve eq. (6) of the main text.

#### A.3.4 Long-lived juveniles

We also present the results for a species with a long-lived juvenile stage (*u*_11_ ≥ 0) but short-lived adults (*u*_22_ *=* 0). Such a life history might reflect that of annual plants with a seed bank, or that of crustaceans like *Daphnia ssp*. with dormant resting eggs. Here, generation time increases with survival in the seedbank *u*_11_.

##### Dispersal and selection only in adults

When selection and migration happen only in adults (i.e. 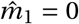 and 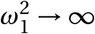), that is in the short-lived stage, we find that genetic and phenotypic divergence becomes

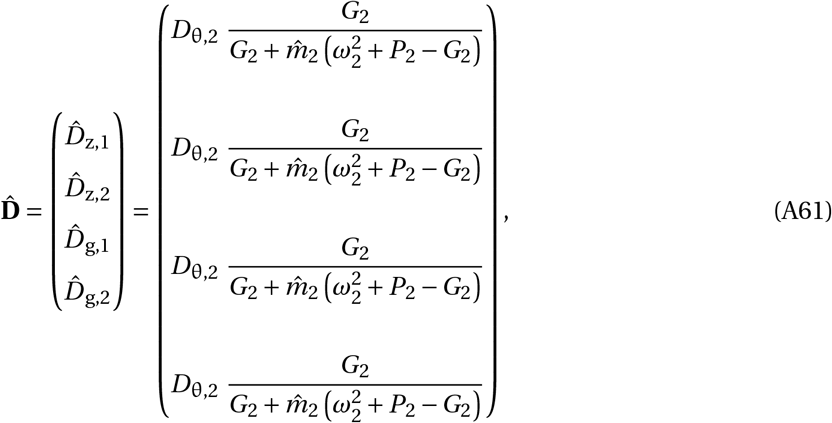

and recover eq. (3) of the main text (at least when ignoring the differences in the stage identifier) for both genetic and phenotypic divergence.

##### Adults under selection and juveniles disperse

With juvenile dispersal and adult exposure to selection (i.e. 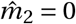 and 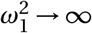),genetic and phenotypic divergence becomes

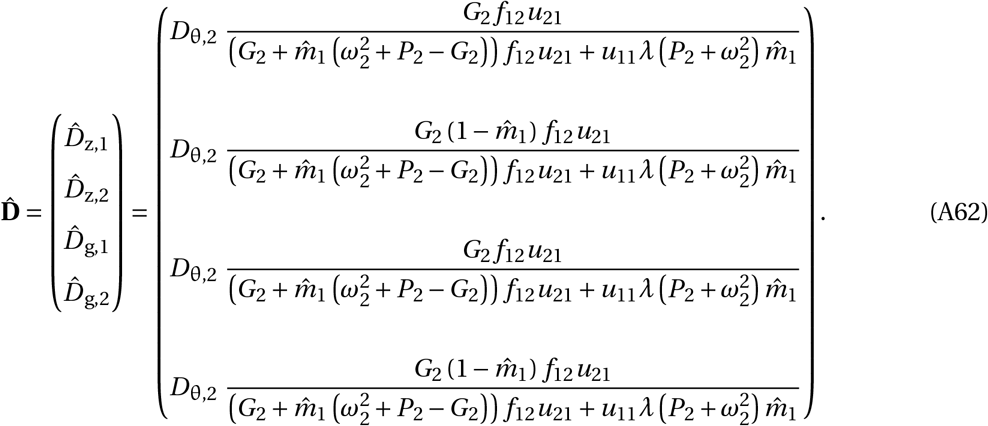

In this scenario, local adaptation declines with increasing juvenile survival *u*_11_ because long-lived juveniles disperse such that dispersal is enforced relative to selection. This result is very similar to the scenario of long-lived adults when selection applies to juveniles but adults disperse (eq. A58).

##### Juvenile selection and adult dispersal

When selection acts on juveniles, while adults disperse (i.e. 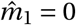 and 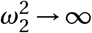), we obtain

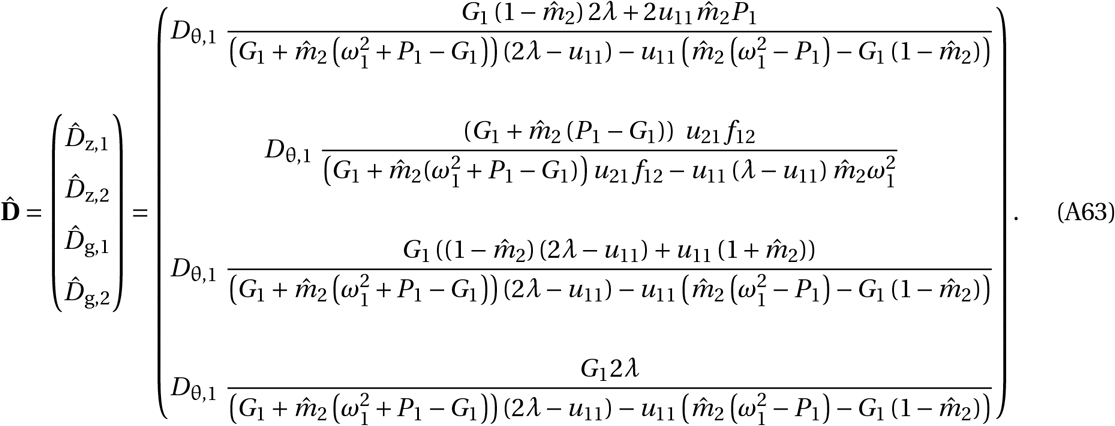

In this scenario, local adaptation increases with increasing juvenile survival *u*_11_ because long-lived juveniles are exposed to selection such that selection is enforced relative to dispersal. This result is very similar to the scenario of long-lived adults when selection applies to adults and juveniles disperse (eq. A59).

##### Juvenile dispersal and juvenile selection

When selection and dispersal acts in juveniles only (i.e. 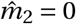 and 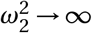), we obtain

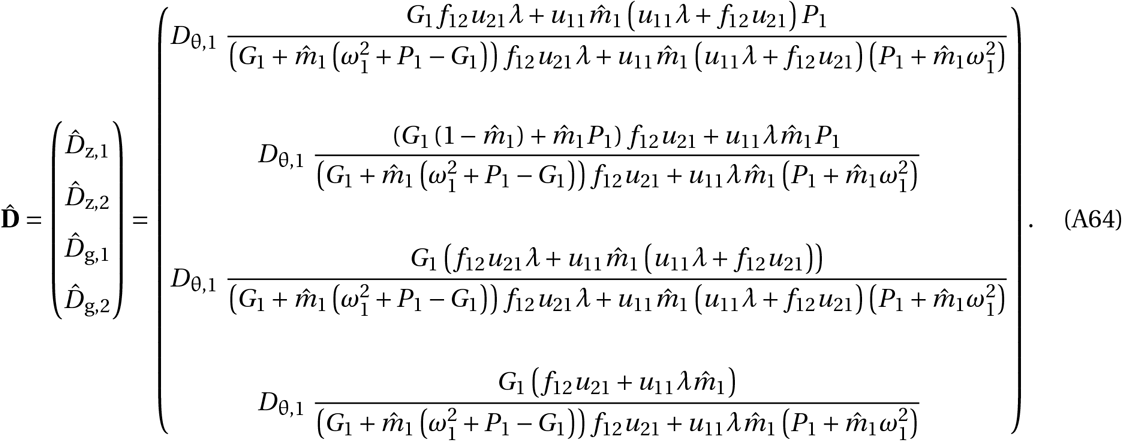

In this scenario, local adaptation decreases with increasing juvenile survival *u*_11_ only slightly because selection and dispersal take place in lockstep. This result is very similar to the scenario of long-lived adults when selection and dispersal apply to adults (see eq. A60).

## B Individual-based simulations

### B.1 Quantitative trait and genetic basis of adaptation

Local adaptation is modelled through a single quantitative trait in diploid individuals. The quantitative trait *z*_*i,s,p*_ is controlled by *n =* 50 loci with additive genetic effects. The trait value is determined by adding up the allelic values at a total of 50 unlinked loci at both homologous copies, using a continuum-of-allele model. Mutations appear with probability *µ*_m_ *=* 0.0001, and mutational effects are drawn from a normal distribution (with a mean of zero and variance *v =* 0.25) and added to the existing allelic value. The total mutational variance *V*_*M*_ then is 2 *n µ*_m_ *v =* 0.0025. Additionally, we model 20 neutral diallelic loci to compare the divergence of quantitative and neutral loci.

### B.2 Life Cycle Events

The sequence of Life Cycle Events (LCE) describes how different processes are handled throughout the simulation (Fig. B1). LCE are stage-specific, when for instance during the reproduction (step 3 in Fig. B1) only adults reproduce. Note that the sequence of LCEs in the simulations follows Fig. 1b, with the exception that life-history transitions are split up into “reproductive transitions” (*a*_12_) and “direct transitions” (*a*_21_, *a*_22_), and that we added a population regulation step to maintain finite populations. Population regulation operates by ceiling regulation when randomly removing individuals (independent of stage or fitness) until the population reached carrying capacity.

**Figure B1:**
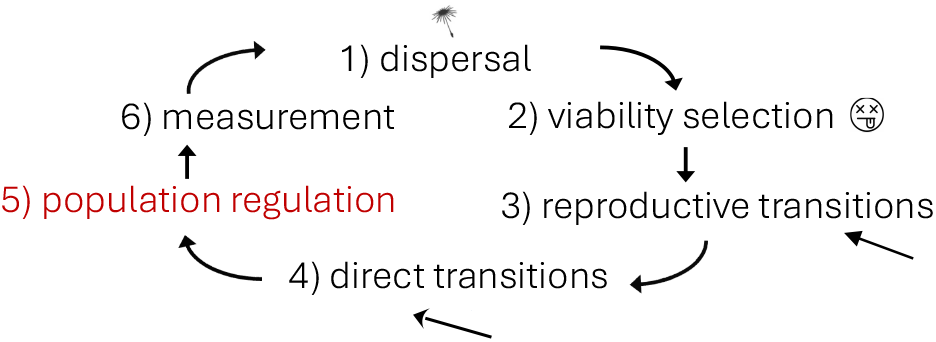
Sequence of LCE implemented in the simulations. The main difference to the yearly events laid out in Fig. 1 is that populations undergo regulation to carrying capacity (highlighted in red).

### B.3 Life-history transitions

As explained in section 2.1, different life history strategies are defined by their matrix population model **A**, where each entry *a*_*sr*_ describes the average contribution of each stage *r* individual to stage *s*. For the simulations, we constructed population matrices

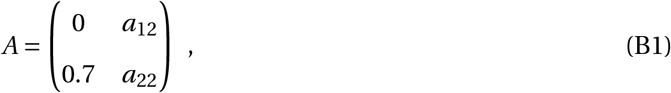

with juvenile maturation *a*_21_ *=* 0.7 (direct transition), with adult survival *a*_22_ (direct transition), and adult fecundity *a*_12_ (reproductive transition). We used adult survival rates *a*_22_ *=* 0, 0.3, 0.6, 0.9, 0.99, 0.999 to manipulate the species’ generation times. To avoid differences in divergence patterns caused by inherent biases in demographic properties, the matrices were standardized for the same population growth rate *λ =* 1.2 (the dominant eigenvalue of the transition matrix (B1)) by adjusting the fecundity *a*_12_ (*a*_12_ *=* 2.06, 1.54, 1.03, 0.51, 0.36, 0.34). Note that this population growth rate is valid for when the population did not reach the carrying capacity yet, and direct and reproductive transitions are the only demographic processes (steps 3-4 in Fig. B1). At demographic equilibrium, however, when the population reached the carrying capacity, we also need to account for population regulation (step 5 of the life cycle in Fig. B1). The “true” vital rates during all life-history transitions (steps 3-5 in Fig. B1) then are the entries of matrix **A** (eq. B1) divided by *λ* (that is *a*_*s,r*_ /1.2).

### B.4 Simulations

Before we extract population statistics, we run burn-in simulations for 50,000 generations at high population size (10,000 individuals per patch, *θ*_*s*,1_ *= θ*_*s*,2_) to achieve selection-migration-balance. After the burn-ins, we run our simulations for further 50,000 generations with *θ*_*s*,1_ ≠ *θ*_*s*,2_ and track genetic divergence 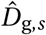,phenotypic divergence 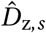,effective dispersal 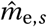,genetic variance *G*_*s*_, phenotypic variance *P*_*s*_, population sizes and stage distributions. We run 10 replicates for each scenario and calculate summary statistics for each replicate. We run two additional sets of simulations, one to assess *G*_*s*_ and *P*_*s*_ before selection (50,000 generations after burn-in, 10,000 individuals per patch) and one to calculate *F*_STQ_ for the quantitative loci and *F*_ST_ for the neutral loci (5,000 generations after burn-in, 10,000 individuals per patch). We run simulations with *D*_θ_ *=* 1, 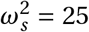, *m*_*s*_ *=* 0.02. All simulations are performed with Nemo-Age (Cotto et al., 2020).

### B.5 Analyses

To assess the evolution of local adaptation, we calculate genetic divergence 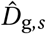 as the mean difference in genotypic trait values between individuals of stage *s* between patches. Besides genotypic and phenotypic trait values to calculate divergence between patches, we also track effective dispersal 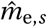,genetic variance *G*_*s*_, phenotypic variance *P*_*s*_, *F*_ST_ and *F*_STQ_. Effective dispersal 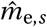 is calculated as the sum of the proportion of immigrant parents among juveniles in each patch, i.e. backward migration. If there are no juveniles present in the population at the end of the simulation round (e.g. when *a*_22_ *=* 0), we calculate the sum of the proportion of immigrant parents among adults in each patch. *G*_*s*_ is measured before selection in the stage under selection. This is done using a separate set of simulations, which allows us to output population statistics before selection. In the absence of plasticity or environmental variance, we assume that *P*_*s*_ is equal to *G*_*s*_. Otherwise, *P*_*s*_ is assessed separately before selection.

All analyses of simulation results were performed in *R* (version 4.2.2; R Core Team, 2022). Graphs were generated using *ggplot2* (version 3.4.0; Wickham, 2016) and *F*_ST_ and *F*_STQ_ were calculated using the package *hierfstat* (version 0.5-11; Goudet and Jombart, 2022).

#### B.5.1 Comparison to mathematical model

In order to compare our simulation results with our analytical model, we calculate analytical predictions for our simulated populations. We calculate the predictions using eqs. (3)-(6), for each respective life-history scenario. The parameters we need for the analytical predictions are either fixed (such as yearly 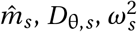 and the “true” vital rates *a*_*i j*_ /1.2 from eq. B1) or need to be extracted from our simulations (genetic variance *G* and phenotypic variance *P*).

### B.6 Results

#### B.6.1 Measuring local adaptation

Throughout the main text, we measure local adaptation as genetic divergence 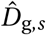 between individuals in two patches adapting to a heterogeneous environment. Alternatively, we can quantify local adaptation in different ways. For example, we can compare the fitness of locals and migrants, as we define local adaptation as granting locals higher fitness in their native habitat than migrants from other habitats. One way to represent this is by comparing survival probability *W*_*s,p*_ (*z*_*i,s,p*_) defined by eq. 1 between locals in a patch and incoming migrants before selection (Fig. B2a). Assessing levels of local adaptation based on local fitness relative to migrant fitness yields the same patterns of local adaptation for the different life histories as observed when comparing 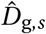 (Fig. 4), even though differences in fitness between locals and migrants are small due to weak selection.

Additionally, we take a closer look at the quantitative loci underlying local adaptation. Firstly, we investigate the covariance of quantitative loci between patches *cov*_*b*_ (see Schmid and Guillaume (2017) Supp. 10), and then we compute the fixation index *F*_STQ_ at quantitative loci (Fig. B2c) and compare *F*_STQ_ to the fixation index *F*_ST_ of 20 neutral diallelic loci in the population, revealing the same patterns of local adaptation once again (Fig. B3).

**Figure B2:**
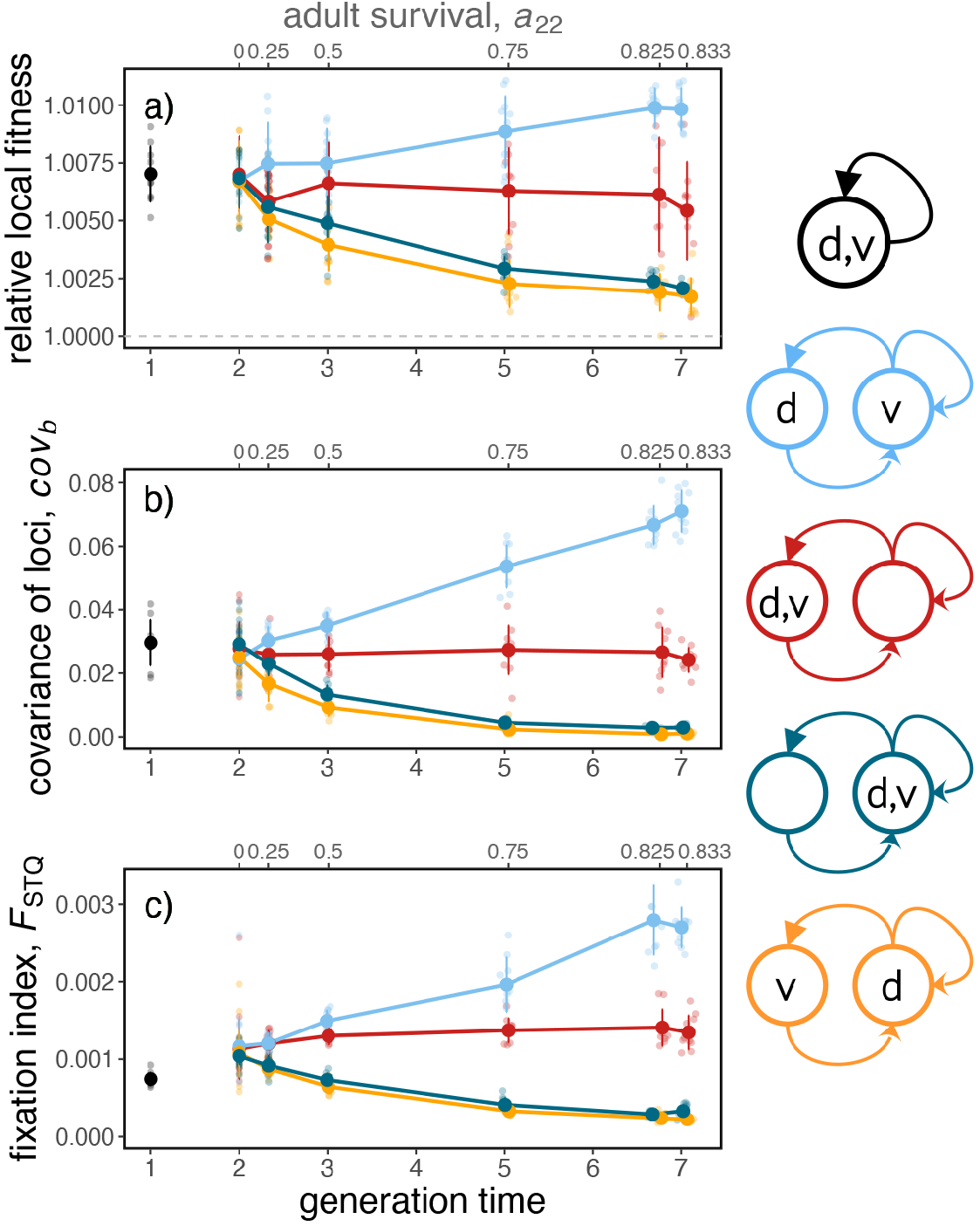
Three alternative ways of revealing local adaptation. **a)** Representing local adaptation using relative local fitness (survival probability of locals divided by the survival probability of migrants). If fitness is independent of an individual’s origin (local or incoming) relative local fitness is equal to 1 (dashed gray line), if locals have higher fitness than migrants relative local fitness is *>* 1, indicating local adaptation. **b)** Looking at the covariance of quantitative loci between patches *cov*_*b*_. Increased *cov*_*b*_ hints at linkage disequilibrium. **c)** Measuring the fixation index *F*_STQ_ for the 50 loci underlying the quantitative trait for adaptation. For comparison of *F*_STQ_ and *F*_ST_, see Fig. B3. All three of these methods yield the same qualitative patterns of local adaptation in the life histories implemented in the main text (compare to Fig. 4). Same parameters as in Fig. 4. Timings of dispersal and viability selection for each species are indicated with letters *d* and *v*, respectively, in life-history diagrams on the right.

**Figure B3:**
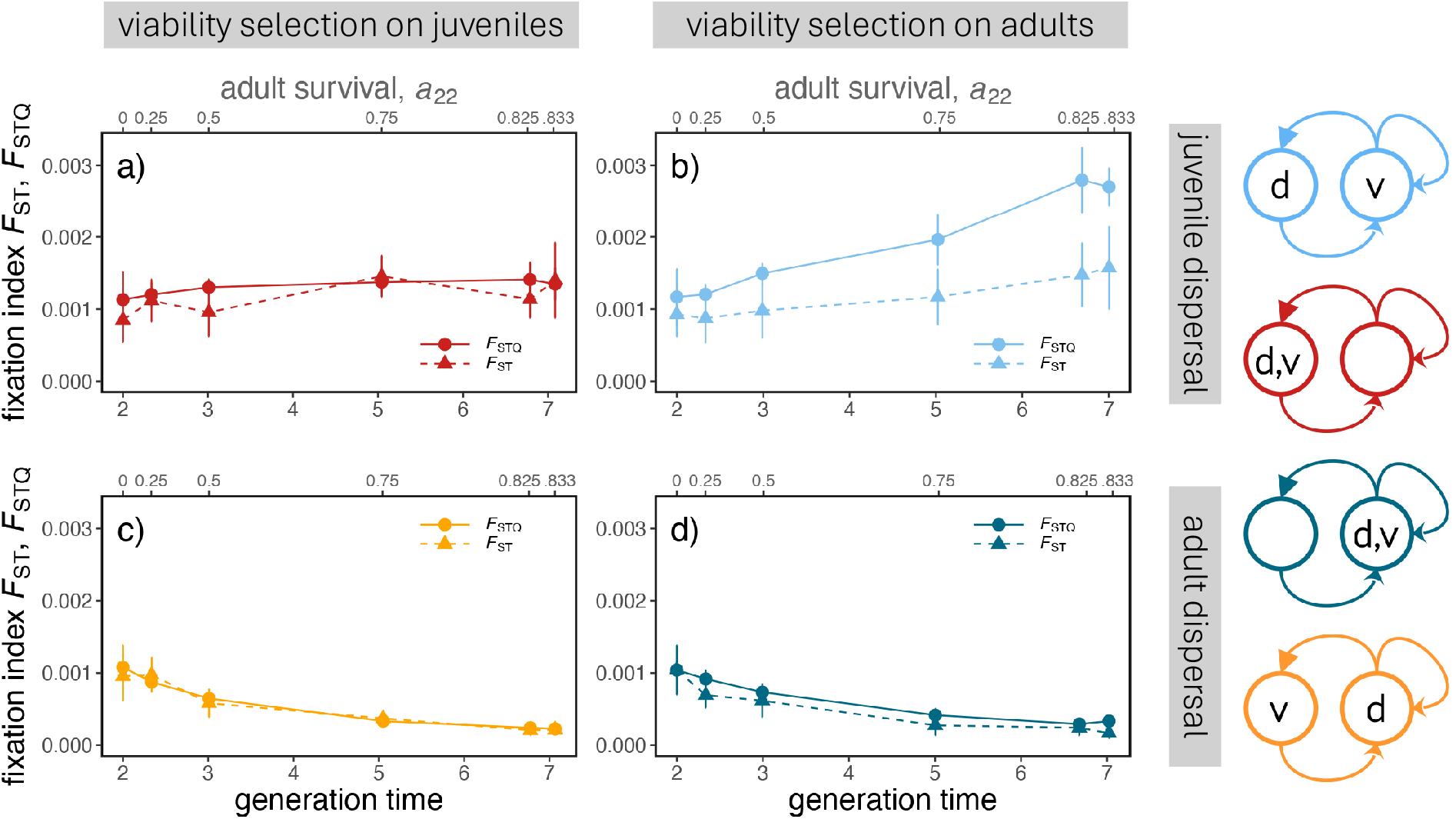
Comparison of neutral and quantitative genetic differentiation using *F*_ST_ and *F*_STQ_, respectively. Increased *F*_STQ_ relative to *F*_ST_ indicates increased genetic differentiation at quantitative loci, indicative of local adaptation. Same parameters as in Fig. 4. Timings of dispersal and viability selection for each species are indicated with letters *d* and *v*, respectively, in the life-history diagrams on the right.

#### B.6.2 Differences between analytical solutions and simulations for long-lived species

We compare our simulation solutions to standard analytical predictions (i.e. for annual species; following equation (A45)) and predictions for long-lived species (following equations (3), (4), (5), or (6), depending on the life history). We calculate the analytical predictions by plugging in the expected yearly forward dispersal 2*m*_*s*_ *=* 0.004 into the mathematical model (Fig. B4). The life history model for long-lived species is able to adjust for increased 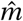 depending on the longevity given by *a*_22_, and thus is able to predict 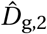 even in life histories where 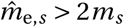.This is especially useful when trying to estimate levels of local adaptation based on natural populations, where it might be easier to assess yearly migration rates *m*_*s*_ rather than effective population mixing 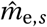.

**Figure B4:**
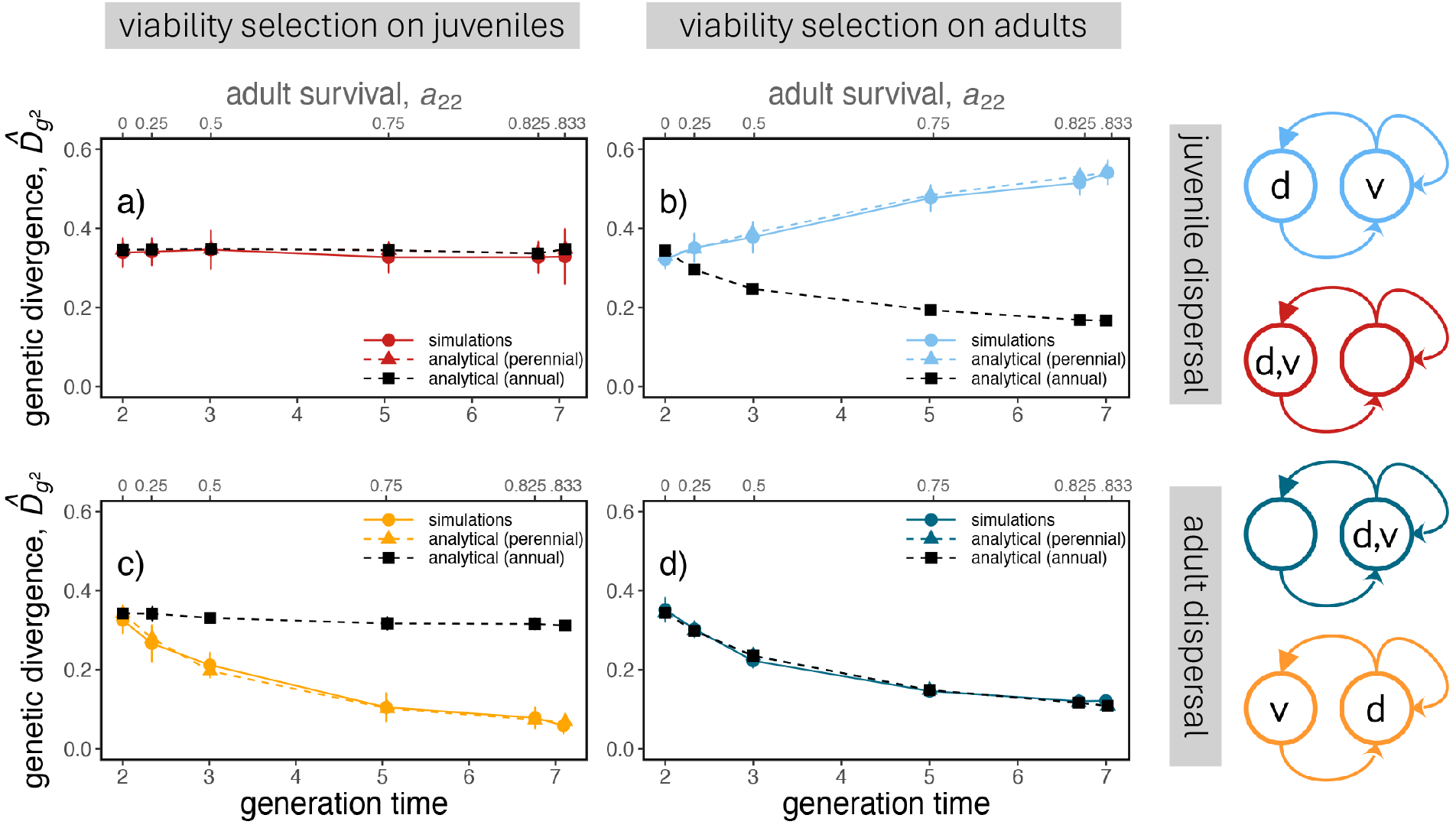
Comparison of adult genetic divergence 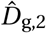 taken from the simulations with analytical predictions based on an annual life history (“analytical (annual)”, eq. 3) and the other life histories elaborated in the main text (“analytical (perennial)”, eq. 4-6). 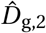 is calculated using the per-year migration rate (forward migration, 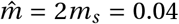). The perennial model matches the simulation results for all life histories, whereas the annual model fails to account for increased lifetime migration caused by longevity in c) (see Fig. 4b for the effect of longevity on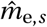) and for strengthened selection in b). Same parameters as in Fig. 4. Timings of dispersal and viability selection for each species are indicated with letters *d* and *v*, respectively, in the life-history diagrams on the right.

#### B.6.3 Difference in phenotypic and genetic divergence

Previously, we assumed there were no environmental effects on phenotypic values for individuals, and genetic divergence was equal to phenotypic divergence. Here, we introduce environmental variance (0.2) to the phenotypic values of individuals. This introduces a slight mismatch between phenotypic and genetic divergence 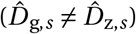 in long-lived species.

**Figure B5:**
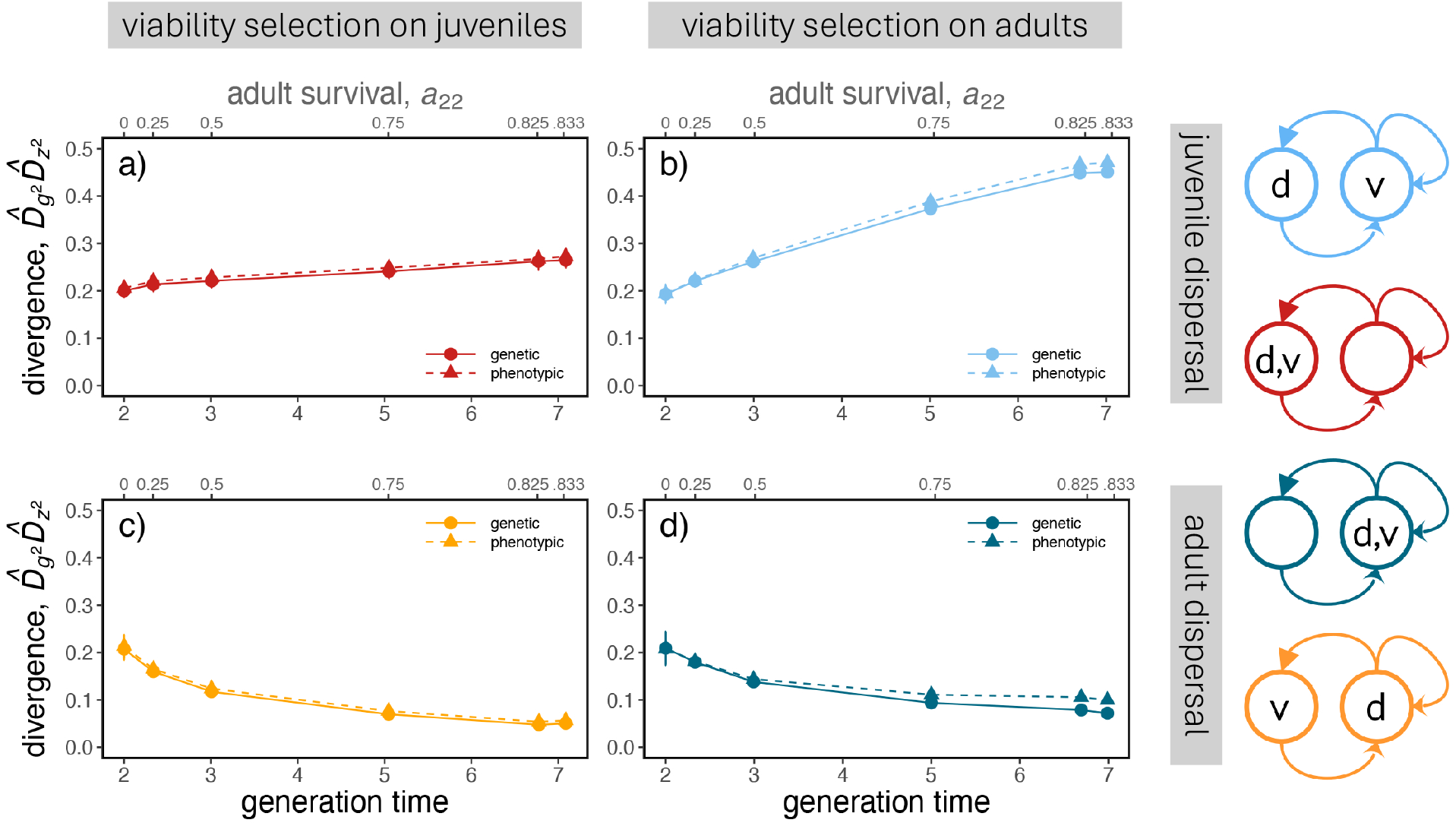
We compare phenotypic (dashed) and genetic (solid) divergence in adults. Phenotypic divergence is slightly higher than genetic divergence in long-living species where adults are under selection b) and d) when there is environmental variance. Parameters are: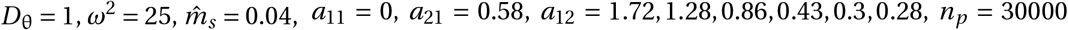30000, 20 quantitative loci. Timings of dispersal and viability selection for each species are indicated with letters *d* and *v*, respectively, in the life-history diagrams on the right.

## Notes

### Competing Interest Statement

The authors have declared no competing interest.

### Summary of Updates

Adding results for a novel life cycle (long-lived juveniles and short-lived adults), improving the figures, adding some more references, and clarifying a section in the Discussion.

